# On Complexity in Resource-Constrained Neuronal Systems: Dynamic Resource Theory

**DOI:** 10.64898/2026.05.20.726716

**Authors:** Kyle Cahill, Mukesh Dhamala

## Abstract

Understanding how complex systems self-organize, exhibit emergent properties beyond their constituent elements remains a challenge across physics, biology, and cognitive science. In resource-constrained neuronal systems, existing theoretical approaches, including gauge theoretic formulations, statistical physics-inspired methods, dynamical population models, and variational principles such as the Free Energy Principle, address important aspects of this problem but do not fully specify the physical conditions and thermodynamic costs under which self-organizing behavior occurs.

Here, we introduce Dynamic Resource Theory (DRT) as a general physical framework for describing self-organization under constrained resource availability. DRT formalizes complexity as a physical property of self-organizing systems arising from coupled mechanisms of resource allocation and dynamic reallocation of internal resources. This framework provides a thermodynamic and variational account of how stability is preserved while adaptive reconfiguration remains possible, consistent with stationary action and thermodynamic constraints. DRT is formulated within a gauge theoretic setting and directly incorporates the energetic costs associated with maintaining structure and enabling system-level reconfiguration.

Within DRT, baseline resource allocation preserves system stability, while internal and external demands perturb the system, driving self-organization through dynamic resource reallocation across a coupled free energy landscape without assuming subsystem separability. We then develop Neural Resource Theory (NRT) and Cognitive Resource Theory (CRT) as principled specializations of DRT, illustrating how this structure is instantiated in resource constrained neuronal and cognitive systems. We conclude by discussing the broader implications of DRT for understanding how complexity, emergence, and adaptive capacity arise over time through thermodynamically permissible reallocation processes across scales.

## 1. Introduction

Complex systems are systems composed of many interacting components and have been widely observed across empirical sciences and extensively modeled across formal sciences [1, 2]. While operational definitions of complexity exist within specific theoretical frameworks, these definitions are typically context dependent and tailored to particular representations or classes of systems [3, 4]. As a result, complex systems are most often characterized in terms of recurring features that arise from interactions among components, including self-organization over time, the emergence of properties, the formation of new structures across scales, and adaptive behavior. For example, Differential Configurational Complexity has recently been introduced as a quantitative measure of spatial organization in scalar field theories, providing a rigorous characterization of localized structure within physically admissible models [5].

Recent work has increasingly emphasized that the behavior of complex systems cannot be fully understood independently of scale, coarse graining, and representation [6]. In many settings, macroscopic or mesoscopic descriptions can be more explanatory or causally informative than microscopic ones, particularly when the phenomena of interest involve coordination, stability, or adaptation [7, 8]. In such cases, the choice of representation can substantially influence which mechanisms are emphasized, as causal influence may be concentrated at scales that are not apparent from fine-grained descriptions alone [9]. Additionally, it’s been demonstrated that spatial inhomogeneity and localization can play a decisive role in determining system behavior, even when coarse-grained or averaged descriptions give the appearance of homogeneity [10].

Across scientific disciplines, complexity is addressed through a diverse collection of modeling paradigms. In neuroscience, entropy-based measures are commonly used to quantify variability and uncertainty in neural activity, dynamical complexity metrics aim to capture the richness and flexibility of temporal dynamics, and criticality-based arguments emphasize proximity to phase transitions as a potential source of adaptive capacity [11-13]. Information-theoretic approaches further frame neural systems in terms of encoding, transmission, and transformation of information [8, 14, 15].

In variational and information-theoretic representations, model complexity and adaptive behavior are often interpreted in terms of variational free energy minimizing criteria [16-18]. Variational free energy minimization is constrained by thermodynamic bounds, since information erasure in physical systems constitutes an irreversible process and incurs at least a minimal thermodynamic cost. This cost, given by the Landauer limit (*k*_*B*_*T* In 2), constrains the set of physically accessible configurations, establishes a fundamental lower bound on the energetics of information processing, and specifies the minimum energy required to irreversibly erase one bit of information [19]. In biological systems, this cost is estimated to be several orders of magnitude above the classical lower bound, with cellular processes operating ∼ 10^1^ − 10^2^ above the Landauer limit [20], which is exceptionally efficient for a physical system. Neuronal computation operates at substantially higher energetic cost, with estimates on the order of ∼ 10^8^ times the Landauer limit, and with communication costs approximately 35-fold greater than computation [21].

There has been sustained interest in grounding neural computation and communication in physical constraints, including energetic costs, metabolic efficiency, thermodynamic, and variational considerations [22, 23]. These efforts have been essential in establishing how neural processing is energetically constrained and metabolically supported [24]. In particular, studies relating metabolic cost, Helmholtz free energy, and variational inference have shown that minimization of variational free energy can provide a sufficient account of both statistical efficiency and metabolic efficiency [22]. From the perspective of a general physics of self-organizing systems, however, a formal framework that makes the thermodynamic costs associated with inference and adaptation explicit across scales has remained an area of active investigation [23].

We demonstrate that a minimal physical description of complex systems can be obtained by clearly describing the energetic constraints that govern system dynamics. We distinguish between two complementary components. The first describes how energetic resources that are allocated to a system governs baseline support, capacity, and constraints across system constituents. The second is resource reallocation, which governs dynamic redistribution of energetic resources in response to demands on the system. Together, these components define Dynamic Resource Theory (DRT), a theory of self-organizing systems from the perspective of foundational level physics intended to formalize how structured behavior can emerge, persist, and adapt under energetic constraints. The relevance of such an organizing perspective is especially clear in neuronal systems, where complexity is well established across spatial and temporal scales [11, 15].

Neural dynamics have been studied across multiple levels of description, ranging from locally coupled Hodgkin-Huxley models [25] to collective population-level descriptions, such as Hopfield networks [26] and other statistical-physics–inspired models. A complementary population-level dynamical tradition emphasizes coordination arising from interactions between excitatory and inhibitory activity, formalized through neural mass models (NMMs) [27, 28].

Phase-based neural mass formulations, including the Kuramoto model and its time-delayed extensions, further demonstrate how coherent behavior and abrupt transitions can emerge from coupled neural populations without centralized control [29-31]. Extensions of this coordination framework to motor behavior, most notably the Haken–Kelso–Bunz model, illustrate how similar dynamical principles govern neural–motor coordination across levels of description [32, 33]

Neural field theoretic models have since extended NMMs into a continuous mean-field description [34-37] which provide principled mesoscopic descriptions of collective macroscopic activity. Statistical physics–inspired methods, including maximum entropy models and related formulations, offer complementary tools that connect collective neural activity to constrained distributions and effective structure for neural interaction which also can be reframed into a variational free energy minimization problem [38-40]. Additional physically motivated perspectives, including gauge-theoretic formulations, further reflect ongoing efforts to describe neural self-organization within principled mathematical frameworks (Sengupta et al., 2016). Across these approaches, however, processes resembling energetic support and redistribution are typically treated as background conditions, inferred consequences, or task-dependent effects, rather than as foundational organizing principles.

To connect this general framework to neuronal systems, we introduce two instantiations of DRT. Neural Resource Theory (NRT) applies the allocation and reallocation structure to living neuronal systems, where resources correspond to biologically constrained energetic and signaling capacities, and reallocation governs coordination, plasticity, and adaptive reconfiguration.

Cognitive Resource Theory (CRT) further specializes this structure to cognition, where task demands impose constraints and where differences in performance may be interpreted in terms of differences in energetic support and reallocation cost, which provides a principled lens for re-examining variability in behavior, efficiency, and adaptation in neuronal networks across individuals and conditions [41].

The structure of this manuscript is as follows. Section 2 surveys major theoretical frameworks that have addressed complexity in neuronal systems, where “major” refers to theoretical scope and the capacity to subsume diverse modeling approaches. These include gauge-theoretic approaches, statistical physics–inspired methods, dynamical population models, and the Free Energy Principle. Section 3 introduces the core formalism of DRT and defines resource allocation and reallocation as its key components. Section 4 develops NRT as a biologically grounded instantiation of DRT in living neural systems. Section 5 extends the framework to cognition by presenting CRT and its interpretation of task engagement, effort, and performance in energetic terms. Section 6 synthesizes the frameworks introduced in Section 2 with NRT, clarifying points of convergence, distinction, and complementarity. Section 7 discusses implications, limitations, and future directions, including opportunities for empirical validation and extensions. Finally, Section 8 concludes by evaluating the DRT framework and clarifying how DRT addresses complexity at the level of foundational physics, situating DRT and its instantiations within the broader study of self-organization in neuronal and other complex systems.

## 2. Existing Frameworks Addressing Complexity in Resource Constrained Neuronal Systems

A central challenge in theoretical neuroscience is to explain how complex, adaptive neural dynamics arise and persist under constraints on energy, information, and physical resources. Neuronal systems operate far from equilibrium, continuously expending energy to maintain functional organization while flexibly responding to internal and external demands.

This section surveys major theoretical frameworks that have addressed complexity in neuronal systems, where “major” refers to theoretical scope and the capacity to subsume diverse modeling approaches. These gauge-theoretic approaches, statistical physics-inspired methods such as maximum entropy models, dynamical population models such as neural mass and neural field formulations, and variational perspectives such as the Free Energy Principle. For each framework, we provide a conceptual overview of its core structure, along with the strengths and limitations intrinsic to adopting it as a basis for modeling neuronal systems.

### 2.1 Gauge-Theoretic Approaches

In physics, a gauge refers to a particular choice of coordinates or potentials used to describe a system whose mathematical formulation contains redundant degrees of freedom. Distinct gauges correspond to different parameterizations of the same underlying physical state, and observable quantities remain invariant under transformations between them. In this sense, a gauge functions as a coordinate system over the space of a field that regulates how energetic quantities are represented, measured, and compared, without altering physical predictions. This invariance reflects the intrinsic symmetry of the system, commonly referred to as gauge symmetry [42-44].

Gauge theory emerged historically from electromagnetism, where different choices of scalar and vector potentials, such as the Coulomb and Lorenz gauges, describe the same electric and magnetic fields. Although these gauges differ in how energy, constraints, and dynamical variables are expressed, they are physically equivalent, and transformations between them leave measurable quantities unchanged. In general, gauge structure provides a formal mechanism for distinguishing physically meaningful invariants from representational redundancy, clarifying which aspects of a model reflect genuine system properties and which arise from coordinate choice. In practice, different gauges may be favored depending on the structure of the problem. For example, in time-independent electrostatic settings, Coulomb or transverse gauges often simplify analysis, whereas in time-dependent regimes involving moving charges, the Lorenz gauge is more natural due to its direct compatibility with special relativity.

These considerations have motivated the application of gauge-theoretic ideas to neuronal systems, where multiple internal descriptions may give rise to equivalent functional behavior. Neural models frequently admit non-unique parameterizations, redundant internal variables, and compensatory transformations that preserve input–output relations or collective dynamics. Examples include degeneracy in neural coding, multiple microscopic realizations of equivalent population-level activity, and alternative dynamical descriptions that yield the same macroscopic behavior [34, 45, 46]. Gauge-theoretic formulations provide a principled framework for organizing such equivalences and for formalizing invariance under reparameterization or local transformation.

Explicit gauge-theoretic approaches to neural dynamics remain relatively sparse, though several works have proposed geometric or symmetry-based formulations to capture invariance, coordination, and constraint in neuronal systems [47, 48]. Notably, the relevance of gauge structure does not depend on scale. In principle, gauge descriptions may be formulated at the level of single neurons, small circuits, or large-scale neural populations.

The relevance of gauge structure in neuronal systems is not limited to explicit gauge-theoretic analysis. The widespread practice of modeling neural dynamics in terms of local potentials, gradients, and field-like quantities already presupposes an underlying gauge structure. Any description that treats differences in electrical potential, current flow, or energetic gradients as physically meaningful, while remaining invariant to an arbitrary frame of reference upon which phenomena such as action potential generation depend, implicitly assumes a gauge symmetry in how those quantities are represented. This is evident even in the most local and biophysically grounded neural models, including the Hodgkin–Huxley formalism and its extensions, where membrane potential, ionic current, and conductance are defined relative to reference frames that do not affect observable dynamics [25, 49-51]. In this sense, gauge structure is not an added abstraction imposed on neural systems, but a latent assumption already embedded in the mathematical language used to describe neural energetics and signal propagation.

While gauge theory provides a powerful framework for organizing symmetry and representational freedom, it is fundamentally concerned with allowable transformations rather than with energetic driving forces. In neuronal systems, gauge structure serves to eliminate redundant degrees of freedom and to provide an effective description of the geometric organization of energetic variables, but it does not specify how such energetic configurations arise. Gauge structure alone does not determine energy dissipation or the mechanisms by which neural configurations are selected, stabilized, or destabilized over time. As a result, additional assumptions about specific physical and biological mechanisms to account for non-equilibrium dynamics and metabolic constraints are essential for describing complexity in resource-constrained neuronal systems.

### 2.2 Statistical Physics-Inspired Methods

Statistical physics offers a principled framework for describing how macroscopic structure and collective behavior emerge from interactions among large numbers of components. Although neuronal systems are open, dissipative, and typically far from equilibrium, equilibrium and near-equilibrium formalisms have proven useful as effective descriptions when the goal is to characterize their statistics, understand collective constraints, and emergent regularities rather than detailed microscopic biophysics [39]. In this sense, statistical physics–inspired models provide coarse-grained representations of neural activity that abstract away detailed dynamics while preserving collective statistics.

A central example of this approach is the maximum entropy (MaxEnt) framework, which constructs the least-biased probability distribution consistent with a specified set of empirical constraints [39]. In practice, one selects a representation of neural state variables such as binary spike indicators, population activity patterns, or discretized functional states. Then one imposes constraints derived from data, including mean activity levels or pairwise correlations. Entropy is then maximized subject to these constraints, yielding a Gibbs distribution whose parameters appear as effective fields and couplings. This construction directly links MaxEnt models to equilibrium statistical mechanics while allowing them to serve as phenomenological descriptions of neural activity [38, 52]

MaxEnt models, despite their name, avoid assuming that neural systems are themselves in thermodynamic equilibrium. Rather, they approximate the stationary statistics of systems that may be dynamically far from equilibrium, treating observed activity patterns as samples from an effective ensemble. In this way, MaxEnt provides a tractable method for capturing nonlinear collective dependencies and higher-order structure in open systems without directly modeling the underlying non-equilibrium processes that generate them [39, 52].

Pairwise MaxEnt models are particularly prominent because they often capture a substantial fraction of collective structure with relatively few parameters, by restricting themselves to first and second order statistics. In these formulations, effective coupling terms are introduced between pairs of units such that the resulting distribution reproduces observed pairwise correlations. Formally, this leads to a model mathematically equivalent to the Ising model, a historically central system in statistical physics originally developed to explain the emergence of collective ferromagnetic order from local spin interactions [53].

The historical significance of the Ising model lies in its demonstration that rich macroscopic structure can arise from simple local interactions, making it a minimal framework for studying collective behavior in neural systems as well [38, 52]. The resulting energy landscape provides a compact representation of likely network states, their relative stability, and transitions between states, implicitly encoding nonlinear interactions among units. In neural populations, weak pairwise correlations can give rise to strongly structured global activity patterns, motivating their widespread use as parsimonious descriptions of collective behavior [38].

These ideas are closely related to energy-based network models, most notably the Hopfield network. Hopfield networks describe associative memory through nonlinear attractor dynamics governed by an effective energy function [26]. Crucially, this energy function acts as a Lyapunov function, guaranteeing that network dynamics monotonically decrease (or leave unchanged) the energy over time, thereby ensuring convergence to stable fixed points (Stern & Shea-Brown, 2020). The existence of such a Lyapunov function depends on symmetry of the weight matrix and the absence of self-connections, linking stability directly to network structure [26]. In this sense, Hopfield networks provide a concrete illustration of how nonlinear dynamics, attractor structure, and stability can emerge from statistical physics–inspired methods.

Statistical physics has also supplied mathematical tools for studying criticality and neuronal avalanches, where neural activity exhibits scale-free distributions and branching-like dynamics. Empirical studies have reported avalanche behavior in cortical recordings, motivating the hypothesis that neural systems may operate near critical points that balance robustness and flexibility [54]. Subsequent work has connected such critical dynamics to large-scale functional organization and resting-state activity patterns, suggesting that statistical signatures of criticality may persist across spatial and temporal scales [55].

Network neuroscience integrates with this statistical perspective by representing neural systems as graphs whose topology constrains dynamics. In this view, network structure defines the space of admissible configurations, while statistical models describe how activity is distributed over that space. Graph-theoretic approaches have therefore been widely used to characterize modularity, integration, and complexity in neural systems, often in conjunction with entropy-based and correlation-based measures derived from statistical physics [56]. Energy-based models and MaxEnt formulations can thus be interpreted as operating over network-defined state spaces, linking structural connectivity to collective statistical organization.

MaxEnt and related statistical physics–inspired models also have successfully been applied to macroscopic neuroimaging data. For example, pairwise MaxEnt formulations have been used to model functional brain states under structural constraints, demonstrating that collective statistical structure persists across measurement scales and modalities [57, 58].

Despite their strengths, statistical physics-inspired models share a set of limitations in their current formulations. Across maximum entropy models, energy-based networks, and criticality-oriented descriptions, the energetic quantities that appear are effective or statistical rather than directly metabolic. The information theoretic structure quantified by these models, such as entropy, surprisal, or correlation structure, can in principle be related to empirical energetic costs using independent physiological constraints. For example, estimates of maximal entropy or information flow may be recast in terms of variational free energy minimization and compared against known energetic expenditures associated with spiking, synaptic transmission, or metabolic consumption [22, 40].

Such correspondences, however, are not specified by the statistical models themselves and instead rely on additional structures outside the model, external measurements, and empirical calibration. As a result, while statistical physics-inspired approaches provide powerful and principled descriptions of collective structure, nonlinear organization, and stability in neural systems, they remain primarily descriptive with respect to the physical energetics that sustain neural activity far from equilibrium [52]. These frameworks therefore characterize collective organization effectively, while leaving open how energetic constraints shape, regulate, and maintain such organization in living neural systems.

### 2.3 Dynamical Population Models

Statistical physics-based approaches to neural organization emphasize collective structure through probabilistic descriptions. In contrast, dynamical population models place emphasis on the time evolution of neural activity, treating nonlinearity, instability, and emergent structure as intrinsic properties of neuronal system dynamics. These models provide a direct mathematical description of how neural populations evolve, interact, and reorganize over time, capturing oscillations, multistability, bifurcations, and spatiotemporal pattern formation that are pervasive in neuronal systems [28, 37, 59].

Dynamical population models represent a predominantly bottom-up, empirically driven approach to modeling neural activity typically at the mesoscale and macroscale. Rather than positing a global organizing principle, these frameworks are constructed from observed features of cortical architecture, population interactions, and collective neural behavior. They are designed to capture dynamics that are difficult to access using local micro-level neuron-by-neuron descriptions, while remaining closely aligned with macroscopic neural measurements such as EEG, MEG, and fMRI [34, 36].

Although dynamical population models are primarily bottom-up in their construction, they are not atheoretical. Neural mass models draw conceptually on the theory of coupled nonlinear oscillators, in which interacting units exchange activity through nonlinear coupling, giving rise to synchronization, rhythmic behavior, and bifurcation structure [28, 60]. Neural field models extend this perspective by adopting a continuum limit, drawing on ideas from classical field theory and mean-field theory to describe the evolution of population activity across space and time (Amari, 1977; Ermentrout, 1998; Bressloff, 2012). In this sense, neural mass and neural field formulations represent complementary limits of a unified population-level description, analogous to the relationship between coupled oscillator systems and spatially extended fields in physics.

A central theoretical foundation of this class of models is the use of coarse-grained approximations, in which large populations of neurons are described by population averaged neural activity. This reduction is motivated by well-established properties of cortical organization, including laminar structure [61], the dominance of pyramidal neurons in long-range coupling and field generation [62], and the substantial redundancy present within local neuronal assemblies. Such redundancy enables principled coarse-graining, whereby microscopic variability is averaged out and collective variables capture dominant modes of activity at mesoscopic scales. As a result, population-level variables acquire physical meaning, and model parameters can be constrained and tuned using empirical observations.

Synchronization is critical in shaping population-level neural dynamics and provides a lens through which collective coordination can be understood [63, 64]. Phase-based descriptions inspired by the Kuramoto model capture how weakly coupled oscillatory populations synchronize through collective interactions, highlighting coherence as an emergent system-level property with excitatory-inhibitory, and time-delayed extensions [29-31, 65, 66]. Related coordination-dynamics frameworks, including the Haken–Kelso–Bunz (HKB) model, emphasize phase transitions and multistability between coordinated modes of activity, providing a principled account of how collective neural behavior reorganizes under changing conditions [32, 33, 67] and coordinates motor response. In related oscillator-based formulations, partial synchronization phenomena such as chimera states, characterized by the coexistence of coherent and incoherent population dynamics, further illustrate how symmetry breaking and multistability can arise in both coupled and uncoupled neuronal networks [68].

Within this framework, neural mass models describe population activity using low-dimensional systems of coupled ordinary differential equations. Canonical formulations such as the Wilson– Cowan equations and their biophysically constrained implementations, such as the Jansen–Rit model, demonstrate how nonlinear excitatory–inhibitory interactions generate oscillations, multistability, and bifurcations, and how variations in coupling strength or external drive qualitatively alter system behavior [28, 69, 70]. These models have been widely applied to the study of rhythmic activity, excitatory–inhibitory balance, and large-scale coordination, and lend themselves to network-based descriptions in which discrete population units are coupled through structured connectivity.

Neural field models generalize these ideas by introducing spatial continuity. Population activity is modeled as a function of both space and time, typically using systems of coupled differential or partial differential equations derived from mean-field limits of densely coupled neuronal networks. Seminal work by Shun’ichi Amari demonstrated that lateral inhibition and spatially structured coupling give rise to pattern formation, traveling waves, and localized activity “bumps” in continuous neural media [37]. Subsequent developments have shown that neural field models can reproduce a wide range of empirically observed phenomena, including wave propagation, spatiotemporal oscillations, and complex attractor structure across cortical tissue [59, 71, 72].

More recently, neural field dynamics have increasingly been analyzed through their spectral and modal structure, in which collective behavior is decomposed into dominant eigenmodes of the underlying field operator or effective connectivity [34, 73]. From this perspective, oscillations, waves, and spatial patterns correspond to the activation and interaction of specific modes whose stability properties govern macroscopic behavior. This eigenmode-based view provides a compact link between anatomical structure, connectivity, and observed neural dynamics, and has been influential in unifying Neural field theory with empirical electrophysiology [35, 74].

Within this spectral framework, multistability and metastability become reducible to a solution set of eigenmodes governing population-level dynamics. Multiple stable or weakly stable attractors may coexist, corresponding to distinct spatial patterns and oscillatory harmonics. Transitions between modes can be driven by changes in coupling, external input, noise, or delays, and have been used to model phenomena ranging from perceptual switching to cognitive flexibility and even pathological transitions such as seizure onset [75-79]. Stability in these systems often reflects higher-dimensional attractor geometry shaped by nonlinearity and temporal structure, rather than simple Lyapunov landscapes, such that population dynamics are organized by multiple interacting modes, delays, and weakly attracting sets that support metastable trajectories and flexible transitions between coordinated states.

Dynamical population models provide a natural setting for describing plasticity and adaptive behavior in neuronal systems. Extensions incorporating activity-dependent changes in coupling or effective connectivity allow neural fields to capture how repeated activation reshapes the dynamical landscape over time. From this perspective, learning and adaptation correspond to slow deformations of the system’s structure, modifying eigenmodes, attractor basins, and stability properties [34, 35, 74].

These models have also been valuable in applied and clinical contexts as well. For example, neural mass and neural field frameworks have been used extensively to study pathological dynamics such as epileptic seizures, where transitions between stable and unstable population-level regimes can be linked to changes in excitation–inhibition balance, coupling strength, or delays [78]. Beyond pathology, eigenmode-based analyses of population dynamics have also been used to characterize distinct functional modes of neural activity, including large-scale coordination patterns associated with cognitive processes such as working memory and task engagement [80]. In this context, shifts between dominant modes reflect changes in population-level coordination rather than localized activation, reinforcing the utility of dynamical population models for linking neural dynamics to cognitive function.

To summarize, dynamical population models of neural activity describe the time evolution of effective population-level variables using continuous fields or low-dimensional state vectors. In neural mass and neural field formulations, collective neural activity is represented by synchronous dynamics governed by coupled differential equations, capturing oscillations, traveling waves, and other emergent phenomena at the mesoscale, as well as their expression at macroscopic scales.

While these formulations specify the temporal evolution of population-level fields, they typically do not provide a closed-form description of an underlying energy operator that specifies the exact physical conditions under which certain dynamics are favored or suppressed under finite energetic availability. Instead, the effective energetic structure of the system is encoded implicitly through empirically informed parameter choices, coupling architectures, and initial conditions. As a result, dynamical population models characterize the space of admissible population dynamics without specifying the energetic mechanism that constrains and biases collective behavior at the mesoscale.

### 2.4 Free Energy Principle

The Free Energy Principle (FEP) formalizes a variational approach to adaptive behavior by applying variational mechanics under a Markov blanket [17, 81]. Under these conditions, the internal states of a system can be shown to follow gradient flows on a variational free energy functional, yielding a mathematically grounded account of how complex adaptive systems such as those found in neural and biological systems maintain their organization and resist dispersion over time.

Within this framework, neural dynamics are interpreted as implementing approximate Bayesian inference under a generative model of the environment [82, 83]. A central structural concept in the FEP is the Markov blanket, which formalizes a statistical boundary between a system’s internal states and external states [84, 85]. By invoking a Markov blanket factorization, the FEP provides a principled definition of system boundaries and specifies that internal states can only infer external causes indirectly, through sensory observations mediated by the blanket.

Under this formulation, internal states are identified with the sufficient statistics of a variational probability density over latent external causes. Specifically, internal states correspond to parameters (e.g., means and precisions) of a variational distribution *q*(*s*) over latent states *s*, while observations *o* denote sensory data accessible through the Markov blanket. The generative model *p*(*o, s*) specifies how observations are expected to arise from latent causes [86, 87].

Given these assumptions, the dynamics of internal states can be expressed as the minimization of a variational free energy functional,

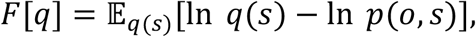

where *q*(*s*) denotes a variational approximation to the Bayesian posterior *p*(*s* ∣ *o*) implied by the generative model. Minimization of this functional corresponds to belief updating that reduces the Kullback–Leibler divergence between posterior and prior beliefs, thereby decreasing prediction error and improving predictive alignment with sensory observations [17, 86, 88].

When applied to biological and neural systems, free energy minimization is typically interpreted from the perspective of active inference, in which systems are modeled as agents that both update internal beliefs and act on the environment to sample informative sensory data [89-91]. In this view, perception, action, and learning reduce uncertainty and thus work in tandem to minimize variational free energy. In this view, processes such as working memory, attention, and salience allocate precision and representational resources under limited capacity [16, 22].

Cognitive effort required to operate these processes is understood as the energetic and informational cost incurred during belief updating [92]. Agency thus emerges naturally from the coupling between inference and control, without requiring additional assumptions beyond the variational formalism.

An important extension of FEP is its formulation in terms of path integrals, which provide an alternative perspective on inference as probabilistic trajectories through state space [17, 93]. Inspired by the path-integral formulation of quantum mechanics [94, 95], this approach represents inference as a sum over possible trajectories weighted by an action functional related to variational free energy. Path-integral representations are particularly useful for analyzing stochastic dynamics, rare events, and long-timescale behavior, and they provide a natural bridge between variational inference, control theory, and statistical physics.

From this variational perspective, many existing models of neural and biological dynamics can be recast as instances of free energy minimization under specific assumptions about generative models and approximations. Statistical physics–inspired models, neural mass models, and neural field formulations can often be interpreted as implementing gradient flows on implicitly or approximately defined variational free energy functionals [40, 96, 97].

Empirical evidence supporting key aspects of FEP has been reported across multiple levels of analysis. At the cellular scale, intracellular recordings in vivo have demonstrated that neuronal membrane potentials and synaptic activity encode prediction errors and precision-weighted updates consistent with variational inference [98, 99]. These findings provide direct physiological support for the interpretation of neural dynamics as approximate Bayesian inference at the level of local neural microcircuits.

At more macroscopic scales, free energy–based models have been shown to account for perception–action cycles and sensorimotor coordination, including eye movements, reaching behavior, and adaptive control under uncertainty, by treating action as a means of minimizing expected free energy through active inference [89, 90, 100]. At the systems level, the Free Energy Principle has been extended to account for large-scale brain organization by framing distributed neural dynamics as processes that minimize expected surprise over extended spatial and temporal scales. In this view, fluctuations in large-scale network dynamics reflect coordinated changes in entropy and uncertainty across distributed neural populations, consistent with predictive models operating at multiple levels of organization [101]. More broadly, the Free Energy Principle has seen applicability beyond neuroscience, and its general formalism suggests a unifying framework for complex adaptive systems [102].

Within FEP, complexity is understood in variational and information-theoretic terms. Specifically, complexity from the perspective of active inference corresponds to the Kullback– Leibler divergence between posterior and prior beliefs under a generative model, quantifying the informational cost of belief updating and maintaining structured internal representations [86, 103, 104]. In this context, degeneracy—where multiple structurally distinct internal states support similar functional outcomes—can enhance adaptability while limiting representational cost, whereas redundancy is reciprocal of efficiency and increases the variational cost associated with added model complexity. From the perspective of active inference, tries to minimize variational free energy by minimizing redundancy while maintaining degeneracy [103].

While Markov blankets define the statistical structure required for inference and allow principled discussions of efficiency and constraint, they do not themselves specify the physical mechanisms that assign energetic costs, dissipation rates, or thermodynamic entropy production; these must instead be established under conditions specified by the generative model [22, 91, 105]. Thus, while FEP provides a clear, powerful, and unifying account of adaptive organization, the extent to which the thermodynamic costs of belief updating and overcoming configurational entropy are compensated by corresponding mechanical benefits remains an active area of investigation [23].

## 3. Dynamic Resource Theory

### 3.1 Introduction to Dynamic Resource Theory (DRT)

In this section, Dynamic Resource Theory (DRT) is formally introduced as a framework for describing the general physics governing complex systems operationally defined by their capacity for self-organization over time and the emergence of structure across scales. The theory is specified by a minimal set of postulates formulated independently of any specific application and grounded in well-established foundational principles of theoretical physics. These postulates assert that the system under consideration is physical, admits a Hamiltonian description of its time evolution, admits a variational formulation consistent with the principle of stationary action, and is subject to thermodynamic constraints generalized to open systems.

DRT is agnostic with respect to the specific dynamical form of the system under study. It does not depend on whether a system is classical or quantum, deterministic or stochastic, linear or nonlinear, open or closed. Rather, what is treated as invariant is the allocation-reallocation mechanism, which specifies the energetic resources that are available to the system and how they are redistributed under thermodynamic and variational constraints. DRT specifies the minimal physical conditions, under the stated postulates, for self-organization to be admissible in systems that admit a Hamiltonian description and obey variational and thermodynamic constraints.

Therefore, it addresses a fundamental question of how organized, adaptive structure can arise at all without violating conservation laws or fundamental thermodynamic constraints.

To establish this framework with maximal conceptual clarity, DRT is developed in the simplest case. The theory is first introduced in a classical, deterministic formulation, then extended to a variational description, and subsequently generalized to the classical stochastic regime. Quantum extensions follow from the canonical quantization of the present formulation. A full operator formalism is reserved as a future direction for this framework and discussed in Section 7.1.3.

#### 3.1.1 Postulates

##### Postulate 1 (Physicality)

A closed dynamic resource system is a dynamic, physical system that evolves in time. As such, it admits a time-dependent Hamiltonian,

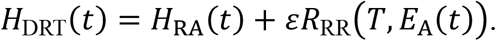

Here, *H*_RA_(*t*) denotes an unperturbed internally regulated configuration of allocated energetic resources to the constituent elements of the system, while *H*_RR_(*t*) denotes a perturbation *εR*_RR_(*t*) that governs thermodynamic resource reallocation.

The quantity *E*_A_(*t*) denotes the available energy for resource reallocation. Here, *T* denotes an effective system temperature defined in the standard thermodynamic sense as the conjugate of entropy,

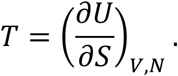

While *T* may evolve on slow timescales due to sustained environmental coupling or internal adaptation, it is treated as quasistatic on the timescales relevant for resource reallocation. The parameter *ε* is the perturbation coefficient quantifying the system’s coupling strength for resource reallocation.

##### Postulate 2 (Stationary Action)

The admissible trajectories of a closed dynamic resource system obey the principle of stationary action,

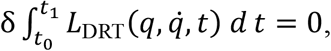

where *L*_DRT_ is the effective Lagrangian associated with *H*_DRT_.

##### Postulate 3 (Thermodynamic Constraint)

Dynamic resource systems in general obey a thermodynamic constraint described by the Jarzynski equality,

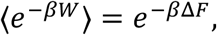

where *W* is the work performed on the system along a trajectory and Δ*F* is the Helmholtz free-energy difference.

In the limit of vanishing work fluctuations ⟨*W*⟩ → *W*, the exponential average collapses and *W* = Δ*F*. More generally, given the convexity of the exponential and Jensen’s inequality,

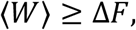

recovers the Second Law of thermodynamics.

### 3.2 Resource Allocation

A dynamic resource system allocates its energetic resources such that the system attains a relatively stable baseline energetic configuration. In general, this configuration corresponds to a steady state that reduces to a dynamic equilibrium in the absence of external driving forces. This allocation reflects the moment-to-moment availability of energy within the system and is shaped by internal fluctuations and structural constraints.

To formalize resource allocation (RA), we introduce a gauge-theoretic description constructed from classical field theory. RA is modeled as an effective field defined over the system’s state space, encoding how energetic resources are distributed across the system’s network architecture.

We define 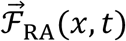 as the negative gradient of a scalar potential Φ_RA_(*x, t*),

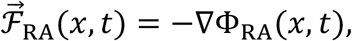

which specifies baseline energetic configuration of the system.

To capture the spatiotemporal transport of allocated resources, we introduce a four-potential 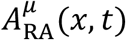, whose components (Φ_RA_, 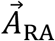) govern the flow of energy across space and time.

From this potential, we construct the associated field strength tensor

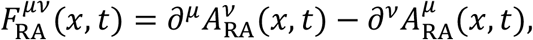

which encodes gradients and circulations in the allocation field.

The gauge freedom of the theory is fixed by imposing the Lorenz gauge condition

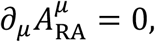

which ensures a well-posed field construction.

The dynamics of the RA field are then specified by the Lagrangian density

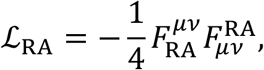

from which the RA Lagrangian is obtained by integrating over three-dimensional space,

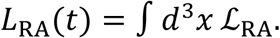

### 3.3 Resource Reallocation

In DRT, Resource Reallocation (RR) is described by a perturbation through which energy is dynamically reallocated when RA is strained. RR reflects the system’s capability to resolve misalignments between allocated resources and energetic demands on the system along the system’s available degrees of freedom, which drives the system’s capacity for self-organization.

This perturbation is defined as

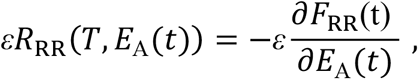

where *T* is the temperature of the DRT system, *E*_A_(*t*) is the energy available for reallocation, and *F*_RR_ is the Helmholtz free energy available for reallocation.

The reallocation function

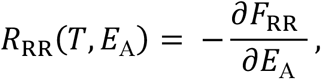

describes gradient descent on the reallocation free-energy along *E*_A_, driving the system toward states that minimize *F*_RR_. RA–RR mechanics is thus governed by a vanishing perturbation. As shown in Figure 1, the system begins at an initial unperturbed allocation of baseline energy (RA), with *H*_RR_ ≈ 0. An event induces resource reallocation (an RR event), introducing a perturbation (+*εR*). This pertrbation transiently modifies the system’s energetic configuration along available degrees of freedom. As the system relaxes, the perturbation is dissipated and absorbed into the baseline allocation, *H*_RR_ → 0. The outcome is an updated resource allocation (RA′) and *H*_RR_ is once again near zero.

**Figure 1.**
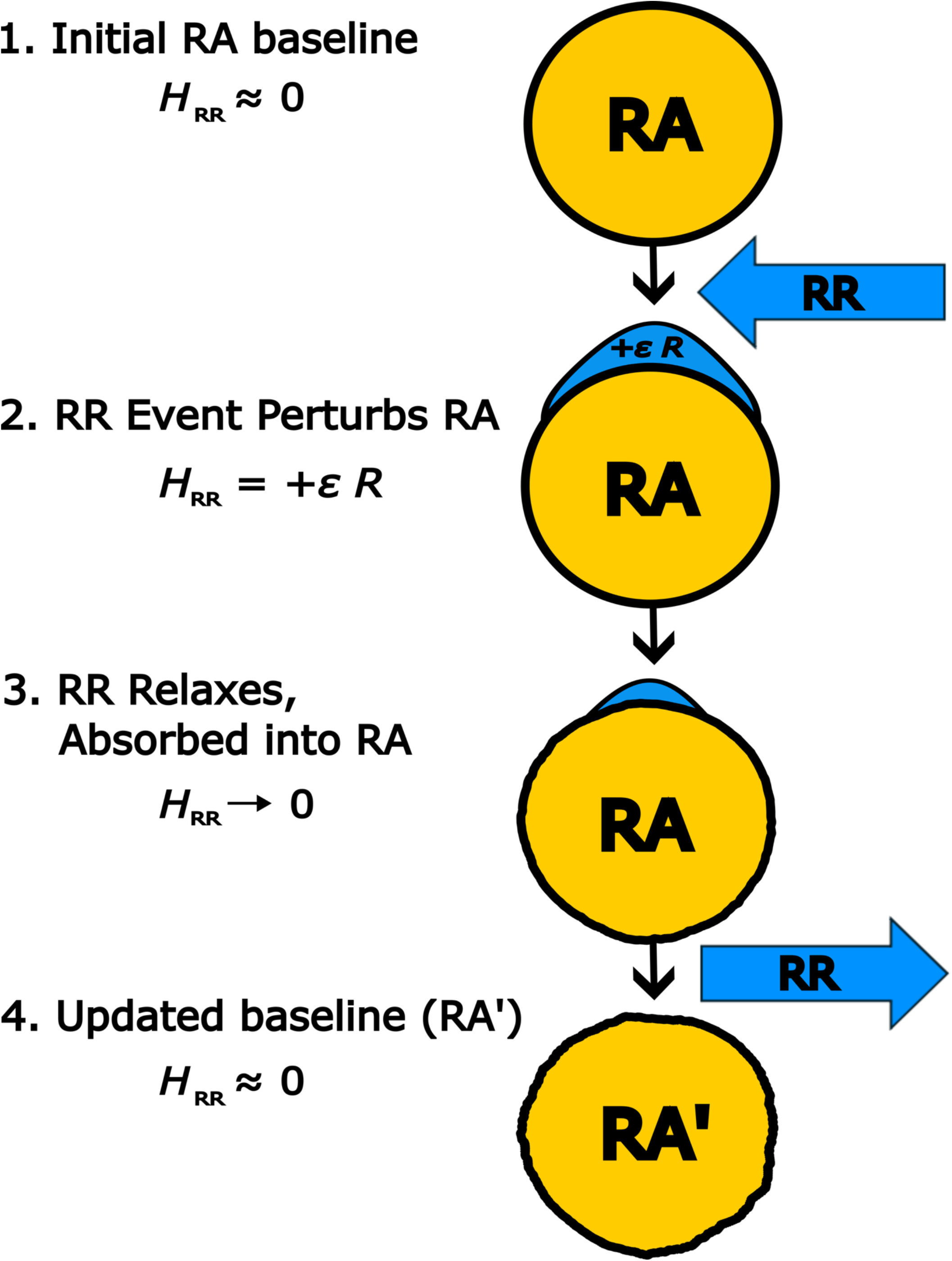
RA–RR Mechanics. Resource reallocation (RR) enters as a perturbation to a system whose energetics is described by resource allocation (RA). An RR event transiently perturbs RA by reallocating resources, corresponding to a modification of the system’s energetic configuration quantified by *H*_RR_. As the system relaxes, the perturbation vanishes and is absorbed into an updated RA (RA’).

#### 3.3.1 Interpretation of *ε*

The parameter *ε* is described as a perturbation coefficient that acts as a coupling parameter that scales the influence of reallocation relative to the stability maintained by RA and reflects the sensitivity of the system’s response to environmental demands or internal fluctuations. When *ε* = 0, the system evolves according to its unperturbed Hamiltonian (RA-only). As *ε* increases, RR exerts proportionally more influence over the system. Typically, *ε* is assumed to be small relative to the unperturbed allocated energy RA, such that *ε*/*E*_RA_ ≪ 1 as in other perturbative systems, reflecting that RR corrections remain small relative to RA.

The perturbation coefficient *ε* scales inversely with the curvature of the reallocation free-energy landscape,

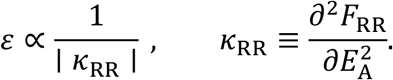

Since *κ*_RR_ has units of inverse energy [J^−1^], the perturbation coefficient *ε* carries units of energy, with *κ*_RR_ quantifying RR stiffness. Although it may initially seem counterintuitive for a perturbative coefficient to have units of energy, *ε* does not represent energy stored in the system. Instead, it reflects the energetic scale set by the inverse curvature of the reallocation landscape. In this sense, *ε* sets the effective gain of reallocation relative to the unperturbed RA dynamics. Small values of *ε* therefore correspond to weak, controlled deviations from the RA-defined baseline, consistent with standard perturbation theory.

We now motivate functional forms for *ε*. Because *E*_A_ and *κ*_RR_ are scalar quantities, admissible parameterizations of *ε* should depend only on scalar combinations of these terms. We therefore consider saturation functions constructed from the dimensionless product *E*_A_*κ*_RR_, which modulates reallocative coupling while preserving energetic consistency.

A general class of smooth parameterizations can be written as

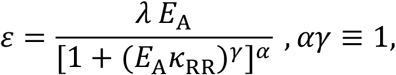

where *λ* is a dimensionless coupling coefficient setting the overall strength of reallocative perturbations, and the constraint *αγ* = 1 enforces dimensional consistency.

In the permissive regime ∣ *E*_A_*κ*_RR_ ∣≪ 1, reallocative coupling is approximately linear, *ε* ≈ *λE*_A_, while in the restrictive regime ∣ *E*_A_*κ*_RR_ ∣≫ 1, reallocative influence is progressively suppressed and asymptotically governed by curvature, *ε* ≈ *λ*/∣ *κ*_RR_ ∣.

Owing to the fact that *ε* carries units of energy, admissible saturation functions must remain real, positive-definite, and bounded for all real values of *E*_A_*κ*_RR_. Because the reallocation curvature *κ*_RR_ may be either positive or negative, physical admissibility requires the saturation function to be even in *E*_A_*κ*_RR_, ensuring sign-independence of reallocative corrections. This restriction limits the saturation exponent *γ* to even integers. Therefore, we adopt the lowest-order, analytically smooth case *γ* = 2, yielding

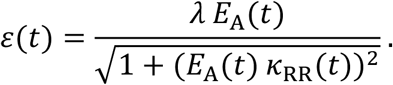

Square-root saturation forms of this type have precedent and are widely used in physics as minimal nonlinear regularizations that ensure effective responses remain bounded while preserving correct low-amplitude limits (e.g., Born–Infeld-type constructions [106]. This choice is also consistent with the perturbative structure of the theory as *ε* represents the lowest-order nontrivial correction.

### 3.4 DRT Thermodynamics

#### 3.4.1 Helmholtz Free Energy in DRT

To determine the Helmholtz free energy of a DRT system, we first define the internal energy of the DRT system. Consistent with standard formulations of analytical and statistical mechanics, internal energy is defined as the expectation value of the system’s Hamiltonian,

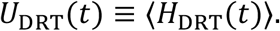

By Postulate 1, the DRT Hamiltonian decomposes into allocation and reallocation contributions,

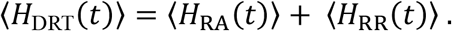

Therefore, the internal energy of the DRT system also decomposes accordingly,

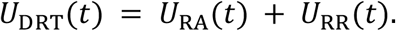

Now we can begin to examine the free energy expressions for DRT’s two core components RA and RR. Since RA defines baseline energetic configuration, its Helmholtz’s free energy *F*_RA_(*t*) is defined as the trade-off between the internal energy *U*_RA_ (*t*), and the thermal energy lost to entropy, given by the product *TS*_RA_(*t*).

This yields the standard expression for Helmholtz free energy,

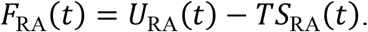

Unlike RA, which corresponds to a stable configuration, RR governs the dynamic response to energetic demands on the system. Because this reallocation process departs from the baseline established by RA, its thermodynamic contribution is structurally distinct and warrants a separate free energy profile.

The free energy of resource reallocation is given by

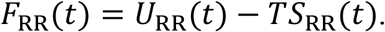

We now define the system’s free energy as the combination of energetic and entropic contributions from both RA and RR. This free energy reflects the system’s state, incorporating both stability and adaptive flexibility. The general form follows the Helmholtz definition

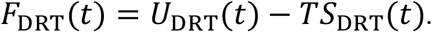

However, we must be careful in describing entropy of a DRT system. Entropy like energy is considered an extensive property, meaning it scales with the size of the system under the assumption that subsystems are independent. For example, doubling the number of independent particles in an ideal gas doubles the entropy. However, for DRT systems RA and RR are not isolated; they share overlapping circuitry, resource channels, and constraints. This means in general for DRT, *S*_DRT_ ≠ *S*_RA_(*t*) + *S*_RR_(*t*), and should instead by calculated holistically using the von Neumann entropy expressed by the system’s density matrix *ρ*_DRT_ which is defined as

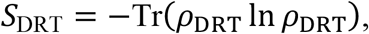

where Tr denotes the trace operator. This formulation captures the system’s total system dispersion under both allocation and reallocation dynamics.

Therefore, *F*_DRT_(*t*), incorporating both RA and RR contributions, is given by

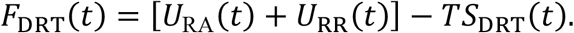

Since DRT is formulated using density matrices and von Neumann entropy, the entropy of the system must account for shared constrained degrees of freedom.

For a state described by density matrix *ρ*_DRT_, with reduced density matrices *ρ*_RA_ and *ρ*_RR_, the entropy decomposition takes the form

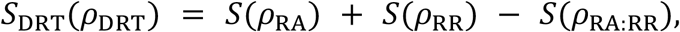

where, *S*(*ρ*) = − Tr(*ρ* ln (*ρ*)) is the von Neumann entropy, and *ρ*_RA:RR_ denotes the density matrix associated with degrees of freedom jointly constrained by allocation and reallocation processes. The subtraction of *S*(*ρ*_RA:RR_) removes double counting of admissible microstates that cannot vary independently due to shared energetic constraints and coordinated dynamics. This correction vanishes only in the limiting case where allocation and reallocation are fully separable.

It follows directly from the entropy decomposition as described above that complexity can be understood as a physical property in self-organizing systems. Specifically, complexity is quantified by the effective entropy of reallocation after accounting for shared entropy arising from degrees of freedom already fixed by baseline allocation,

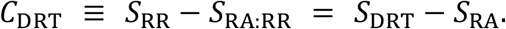

This measure captures the number of physically admissible reconfigurations available to the system under a fixed allocation of internal resources. In this sense, complexity reflects the system’s capacity for adaptive reorganization—the multiplicity of energetically allowed ways in which internal resources may be redistributed without altering the underlying structural constraints imposed by RA.

Substituting this entropy decomposition into the Helmholtz free energy yields

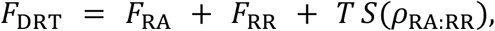

showing that allocation–reallocation coupling introduces a thermodynamic correction to the system’s free energy. This term quantifies the energetic cost or benefit associated with maintaining coordinated resource configurations, providing a principled physical interpretation of allocation–reallocation coupling beyond phenomenological interaction terms. Importantly, the non-negativity of *S*(*ρ*_RA:RR_) guarantees consistency with Postulate 3 of DRT.

#### 3.4.2 Free Energy Minimization in RR

So far, we have assumed the form of the reallocation function as a thermodynamic function described by

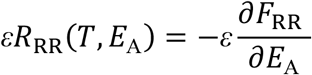

but up to now we have not yet shown proof as to what form *R*_RR_ should necessarily take. Since *R*_RR_ is a physically consistent dynamic reallocation of a system’s energetic resources, it should depend at least on the available energy for reallocation *E*_A_(*t*) and reallocation must occur within some environment which implies a dependence on *T* defined as the effective temperature of the system.

Therefore, *R*_RR_(*T, E*_A_(*t*)) specifies the minimal dependencies required for a reallocative mechanism to operate. The available energy *E*_A_(*t*) denotes the energy available for reallocation which excludes energy allocated by RA, *U*_RR_(*t*) = *E*_RA_(*t*), as well as energy locked (bound) into energetically inaccessible degrees of freedom that are essential for system stability, denoted *E*_L_. The energy available to reallocation is thus given by

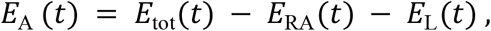

where *E*_tot_(*t*) represents total energy of the system at a given moment in time. In order to describe physically admissible reallocation, it is necessary to formulate a physically consistent expression for the reallocation function *R*_RR_(*T, E*_A_). Since *R*_RR_(*T, E*_A_) is a mechanism that redistributes energy *E*_A_, it should be consistent with thermodynamic constraints specified by Postulate 3, namely, the tendency of physical systems to minimize free energy. In this context, the Helmholtz free energy of RR, *F*_RR_(*t*), is the quantity to be minimized.

To determine how the system minimizes *F*_RR_(*t*), we proceed by taking partial derivative with respect to the available energy, *E*_A_ (*t*),

This yields,

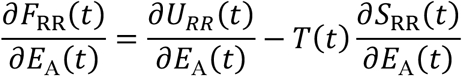

To ensure that reallocation proceeds toward a physically meaningful steady state, we require that the system evolves such that *R*_RR_(*T, E*_A_) → 0, corresponding to a gradient descent of the reallocation free energy *F*_RR_ along *E*_A_.

Under this condition, *R*_RR_ tends toward zero when the magnitude of the free-energy gradient vanishes,

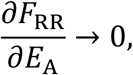

which implies

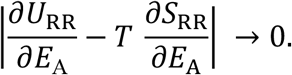

From dimensional analysis, *T, S*_RR_, and *E*_A_ are all positive-definite quantities with consistent physical units. Moreover, since RR constitutes a dynamic subsystem—exchanging energy with other system components—its thermodynamic behavior is necessarily bounded.

For a DRT system in a steady state at temperature *T*, the entropy change per unit of available energy must therefore satisfy

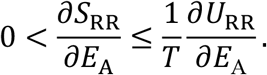

Since *U*_RR_ = *⟨ϵ*(*E*_A_) *R⟩* and *ϵ*(*E*_A_) *i*s monotonic and strictly positive for all *E*_A_ > 0, *U*_RR_ increases with *E*_A_ whenever |*R*_RR_| > 0.

Free-energy minimization would then proceed until the system saturates its capacity for configurational reordering, such that further energetic investment produces no additional increase in *S*_RR_, at which point

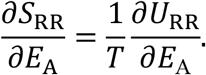

Therefore, the expression that satisfies the minimization criterion is

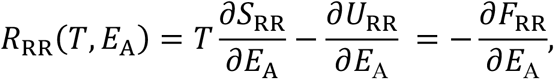

arriving at a final expression for the reallocation function given by

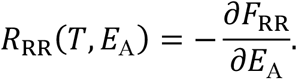

Using the form reallocation function *R*(*E*_A_) and perturbation coefficent epsilion *ε*(*E*_A_) derived in Section 3.3.1, in Figure 2 we demonstrate the resulting RA-RR dynamics under closed system conditions. For illustrative purposes, we approximate the reallocation entropy as

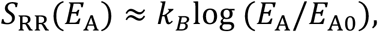

where *E*_A0_ denotes the energy available at the onset of the reallocation event. This approximation reflects the minimal assumption that accessible reallocation configurations scale monotonically with available energy. We adopt a local first-order Taylor expansion of the reallocation internal energy with respect to the available energy, setting 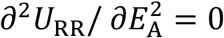. This simplifies the model while preserving the thermodynamic structure of RR.

**Figure 2.**
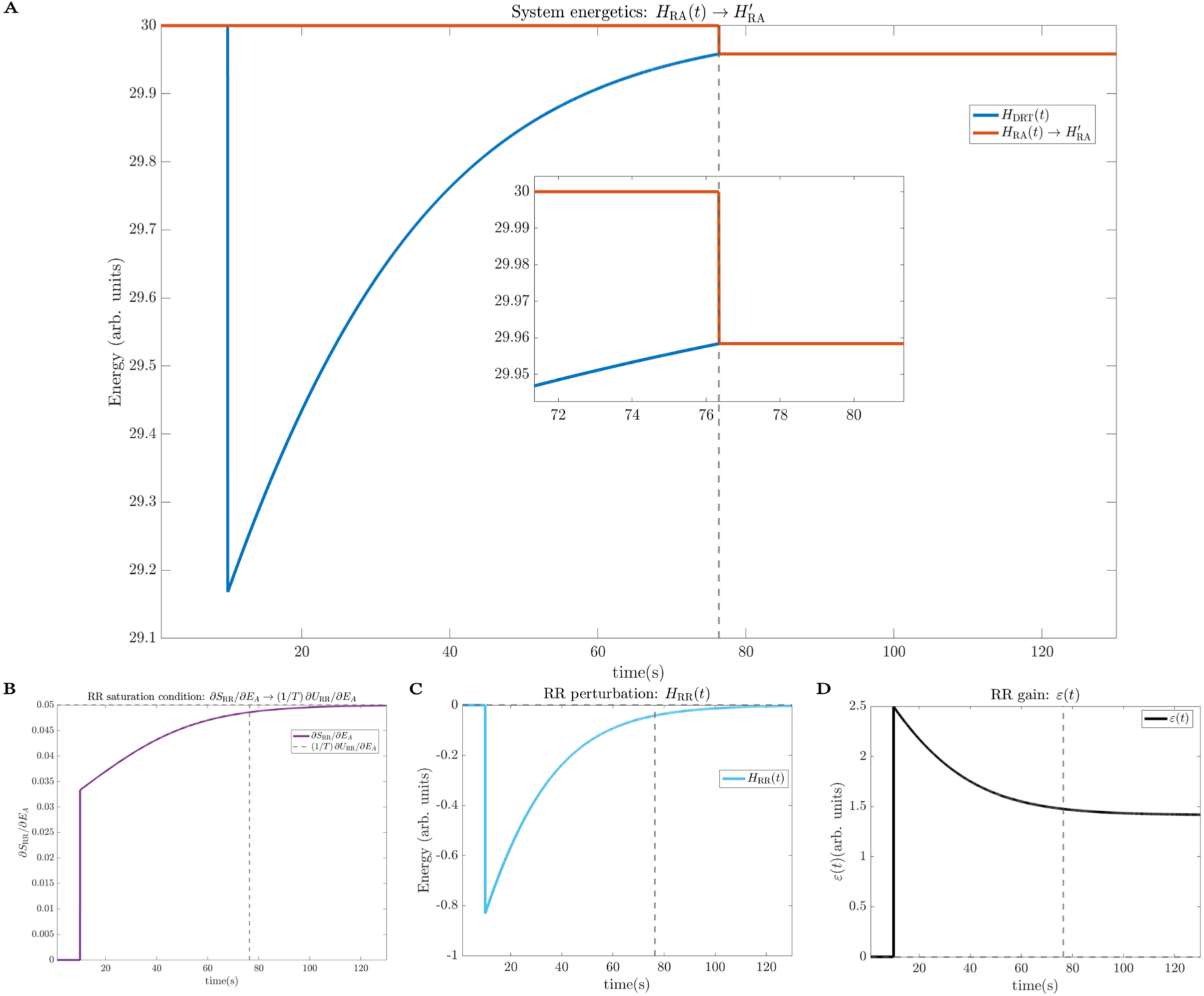
RA–RR dynamics under a closed-system, first order approximation. (A) System energetics showing *H*_DRT_ = *H*_RA_ + *H*_RR_ and the post-reallocation update 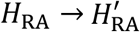. The dashed line marks a practical convergence criterion (5% of its initial perturbation magnitude), below which ∣*H*_RR_∣ becomes effectively indistinguishable from the updated allocation baseline 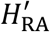. Inset: zoomed view of the convergence. (B) RR perturbation *H*_RR_(*t*), showing decay toward zero. (C) RR perturbation gain *ε*(*t*). (D) RR saturation condition ∂*S*_RR_/ ∂*E*_A_ → (1/*T*) ∂*U*_RR_/ ∂*E*_A_, defining the thermodynamic equilibrium condition for reallocation. Parameters at RR onset: *H*_RA0_ = 30, *E*_A0_ = 30, *T* = 20, *k*_*B*_ = 1, *λ* = 10^−2^. All energies are shown in arbitrary units with *k*_*B*_ = 1; time is expressed in seconds.

#### 3.4.3 DRT Partition Function

At any given moment in time, a DRT system occupies states probabilistically, with each state’s likelihood determined by its energetic cost *E*_*i*_ and an effective temperature *T*. At the level of ordinary matter composed of atoms and molecules, Boltzmann statistics under quasistatic conditions provide the simplest and most effective description of state occupancy. Under these assumptions, the system admits a Boltzmann distribution, defining the probability of occupying a state *i* as

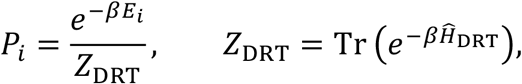

where *β* = 1/(*k*_*B*_*T*), and Tr denotes the trace operation, corresponding to a sum over all accessible system states. In practice, this involves summing the diagonal elements of the matrix representation of the Hamiltonian 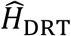, with each diagonal entry corresponding to the energy assigned to a distinct configuration of the system. The Hamiltonian is expressed as an operator (denoted by the hat) to emphasize its role in assigning energies to the space of possible configurations, ensuring proper normalization of the resulting probability distribution.

Here, *P*_*i*_ denotes the probability that the system occupies a state with energy *E*_*i*_, given by the eigenvalue 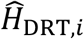 of the DRT Hamiltonian evaluated at state *i*. Higher-energy states are exponentially suppressed, while the normalization constant *Z*_DRT_, known as the partition function, encodes the energetic weight of all accessible states under DRT dynamics. In DRT, the Hamiltonian is constructed as the sum of allocation and dynamic reallocation contributions, consistent with the first law of thermodynamics, 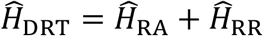. This decomposition reflects the fact that the system’s energy includes both a relatively stable allocation component 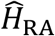 and a dynamic reallocation component 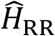.

#### 3.4.4 DRT Entropy

The von Neumann entropy quantifies internal dispersion within a DRT system as

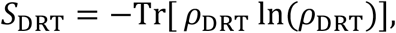

where *ρ*_DRT_ is the system’s density matrix, encoding the probabilistic distribution over states. The challenge, then, is to construct a physically plausible expression for *ρ*_DRT_, the system’s density matrix. To incorporate empirically grounded features such as synchrony, phase-locking, and interregional coordination, which are hallmark examples of system self-organization, RR is modeled using an Ising-like structure.

The Ising model was originally developed in statistical physics to explain ferromagnetism, specifically how simple local interactions between neighboring atomic spins can give rise to large-scale collective magnetic order [53]. In the context of DRT, this same structure offers a principled way to model coordinated reallocation under energetic constraint. In this formulation, RA is diagonal by construction, providing a stable baseline that admits a Hamiltonian description of the system. RR introduces coupling between units via pairwise interaction terms, aligning with empirically observed synchrony, coordination, and emergent dynamics characteristic of self-organizing systems.

The corresponding DRT Hamiltonian is expressed as

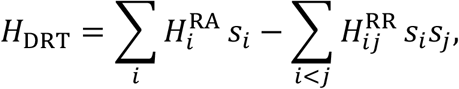

where *s*_*i*_ ∈ (−1, +1) represents whether unit *i* is active (+1)or inactive (−1), following conventions from spin-based models in physics. Here 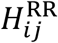 represents the pairwise interaction energy between units *i* and *j*, introduced by RR during episodes of resource reallocation.

The density matrix under this model is expressed as

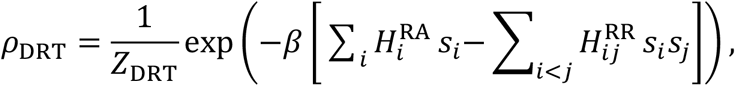

with partition function

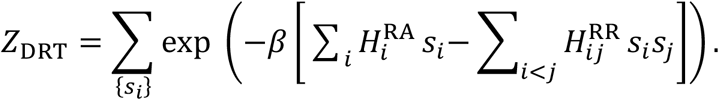

This formulation provides a physically grounded basis for computing entropy in DRT under an interacting, self-organizing model of resource reallocation, with *β* = 1/(*k*_*B*_*T*), where *T* denotes the system temperature and *k*_*B*_ is the Boltzmann constant.

A key feature of DRT is that it admits a quantitative notion of complexity as physical property. In particular, the structure of the reallocation Hamiltonian *H*_RR_ determines the accessibility of possible reconfigurations of internal resources. This multiplicity of accessible reconfigurations is effectively measured by the reallocation entropy. In the absence of accessible reconfigurations, a system is restricted to states admissible solely owing to resource allocation, corresponding to a comparatively simple system with no capacity for self-organization among its internal degrees of freedom.

In the limiting case where the RA and RR Hamiltonians are assumed to commute, each may be modeled as a paramagnet, corresponding to a non-interacting system in which units behave independently. Under this assumption, the entropy *S*_DRT_ becomes additive, with each unit contributing independently to the system’s entropy.

If each unit contributes only local energy terms, this yields a fully diagonal Hamiltonian

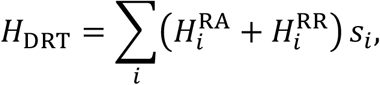

where *s*_*i*_ ∈ (−1, +1).

The corresponding density matrix reduces to

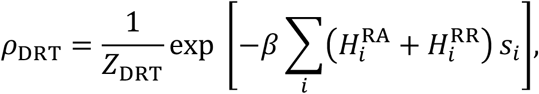

with partition function

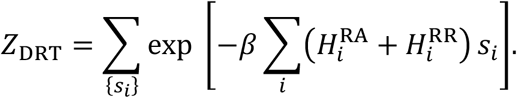

This formulation is computationally efficient and analytically separable, providing a physically density matrix for DRT entropy in the absence of interaction, omitting RA-RR coupling terms.

### 3.5 Variational DRT

#### 3.5.1 Lagrangian Formulation and Variational Action in DRT

Physical systems tend to evolve along paths that minimize a quantity known as action—a measure of the system’s energetic expenditure across time. This principle, formally referred to as the principle of stationary action, forms Postulate 2 of DRT and is more commonly known as the principle of least action, both because minimization is the typical case and because the phrasing is more intuitive. The concept aligns with everyday metaphors like “following the path of least resistance” or “going with the flow”, expressions that echo nature’s preference for efficient trajectories. In physics, this principle provides a powerful framework for understanding how dynamic systems behave under constraints. To capture this formally, DRT defines an action functional, which quantifies how costly a given pattern of reallocation would be. The system then follows the trajectory that minimizes this quantity, yielding the most energetically favorable path given current constraints.

To model DRT systems, we define a Lagrangian for the evolution of state variables given by generalized coordinates *q* which correspond to the degrees of freedom a system’s constituent elements may have. The baseline dynamics are still governed by RA which defines a minimal-energy trajectory. Deviations from this baseline are mitigated by RR which dynamically modulates energy across subsystems. The rate of change 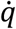 captures how quickly the system’s state changes over time and may correspond to observable dynamics. Together, *q* and 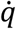 describe the system’s evolving trajectory through state space. This trajectory reflects both the intrinsic dynamics (governed by RA) and demand-evoked adjustments (modulated by RR), defining how systems reconfigure. To formalize these dynamics, we define the DRT Lagrangian as the sum of a baseline term provided by RA and a perturbation term introduced by RR

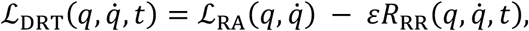

where 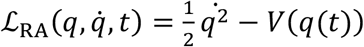 is the RA Lagrangian. The function *V*(*q*(*t*)) represents the potential energy associated with the system’s current configuration.

Although it may formally depend on time, it is standard practice to express this simply as *V*(*q*) with the time dependence understood implicitly. 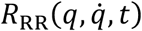 is the perturbation term encoding the demand-induced reallocation function, while *ε* is the perturbation coefficient that quantifies the strength of RR influence (see Section 3.3).

To unify the Lagrangian and Hamiltonian formulations of DRT, we apply the Legendre transform, and we set the effective mass to unity, *m* = 1, to simplify the kinetic term. Starting from the Hamiltonian,

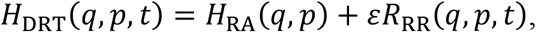

where 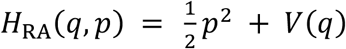. The velocity can be obtained from Hamilton’s equation denoted as, 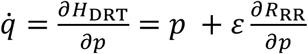.

The Lagrangian is then obtained by the Legendre transform given by

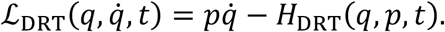

Substituting 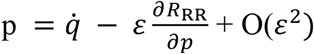, we can then write the Lagrangian as,

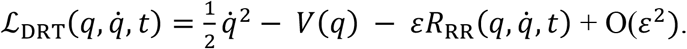

This shows that the reallocation term *R*_RR_, can be meaningfully transformed between Hamiltonian and Lagrangian formalisms and thus validating its appearance in the Hamilton– Jacobi equation given by

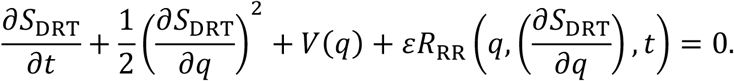

#### 3.5.2 Variational Representation of DRT

In the variational representation of DRT, the system is characterized by possible trajectories through state space weighted by their associated action. These trajectories are governed not only by baseline energetics but also by transient reallocations induced by RR. To capture this, a path integral over histories parameterized by the generalized coordinates *q*(*t*), which describe the system’s degrees of freedom, incorporates both RA and RR components into the action.

This treats adaptation as a superposition of possible trajectories, each weighted by its energetic cost. In the classical limit, the dominant trajectory minimizes the action, producing efficient yet constrained system response. The path integral over *Dq*(*t*) sums over all admissible trajectories.

Operationally this expression is defined as,

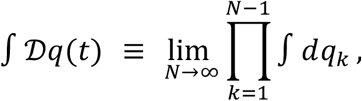

where *q*_*k*_ ≔ *q*(*t*_*k*_) are the system’s configuration variables evaluated at discrete time slices *t*_*k*_, and the endpoints *q*_0_ and *q*_*N*_ are fixed by boundary conditions, with each trajectory weighted by an exponential kernel derived from the DRT action. This kernel reflects the energetic cost of the path, encoding the system’s probabilistic preference for efficient reallocation. To describe such a path integral representation for all possible paths a system might take, and expectation of observable quantities based on these possible paths, following inspiration from Feynman’s path integral formalism [94, 95] we must first describe the DRT partition function *Z* which is denoted by

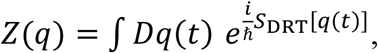

where the DRT action *S*_DRT_[*q*(*t*)] is expressed as

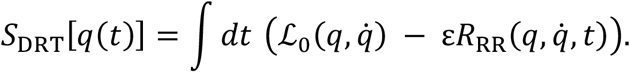

While DRT adopts the formal machinery of action-based dynamics, including path integrals and partition functions, the DRT action *S*_DRT_[*q*(*t*)]is not derived from fundamental fields. Instead, it is a phenomenological effective action defined over system state space, encoding energetic, entropic, and constraint-based structure relevant to self-organization. Establishing a direct correspondence between DRT and field-theoretic Lagrangians would require mapping onto underlying degrees of freedom and is therefore reserved for future work.

In formalizing DRT, we adopt the convention of natural units by setting the reduced Planck’s constant to unity (*ħ* = 1). In this context, this reflects a move to natural units to simplify and generalize the structure of path weighting. By doing so, the action becomes dimensionless, allowing the dynamics to be expressed in terms of relative path weights rather than absolute units. This convention enables the partition function *Z* to take the form

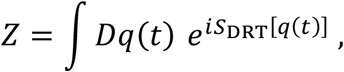

which allows calculation of expected values of observables ⟨*𝒪*[*q*(*t*)]⟩ given by

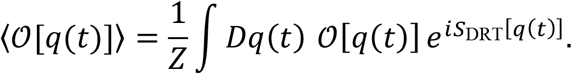

Examples of ⟨*𝒪*[*q*(*t*)]⟩ include *ℰ* which is the expected energetic cost of a trajectory; *ρ*_*ij*_ which is a scalar measure of pairwise synchrony between the system’s constituents, T which is expected temperature across all trajectories and *J*[*q*(*t*)] which describes “negentropy” or rather entropy reduction from a maximal baseline. Negentropy quantifies internal structure or representational precision gained along a trajectory. As an example, how to calculate an expectation value under this formalism, consider the expected energetic cost *ℰ*[*q*(*t*)] of some external demand on the system.

To do this, we write the energy functional as

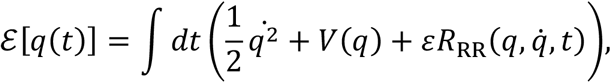

then the expression,

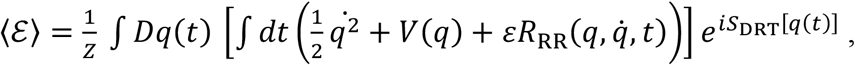

yields the expected energy which quantifies the average cost the system will expend across all feasible paths when responding to an external demand on the system.

Critically, the support of the path integral, which corresponds to trajectories that contribute to the value of an observable, ⟨*O*⟩, is shaped by the structure of *S*_DRT_[*q*(*t*)]. From this foundation, the DRT partition function encodes not only optimal strategies but the full probability-weighted distribution of trajectories that a system could take under real-world constraints. In the absence of strenuous demands, RA governs minimal-effort dynamics along energy-efficient paths. In response to strain RR reallocates energy to meet the system’s expected energetic demands. This shifts the system’s trajectory away into a new region of state space, governed by the modified Lagrangian.

In future extensions, it may be plausible and, in some cases perhaps necessary, to introduce a normalization constant *c*, acting as a global scaling factor over the space of trajectories. In traditional path-integral formulations, such constants are typically absorbed into the normalized partition function *Z*. Within DRT, however, a scaling factor may carry theoretical significance, reflecting effects of coarse-graining, system-specific tuning governed by global resource constraints, or the characteristic scales at which a system exhibits meaningful structure. Although this constant is not modeled here, its inclusion could serve as an effective renormalization parameter, enabling controlled variability across systems or realizations described within the DRT framework.

### 3.6 Stochastic DRT

Having established the core structure of Dynamic Resource Theory in the deterministic and variational settings, its extension to classical stochastic dynamics follows naturally. The allocation-reallocation mechanism central to DRT is invariant across deterministic and stochastic representations.

In stochastic systems, the system state evolves probabilistically rather than along a single trajectory, while the underlying energetic structure remains well-defined. Standard stochastic formalisms such as Langevin and Fokker–Planck descriptions readily accommodate Hamiltonian structure and thermodynamic constraints. The same allocation–reallocation structure that governs deterministic dynamics continues to apply in the stochastic regime, now expressed through probabilistic transport rather than single trajectories.

While openness and stochasticity are independent properties of dynamical systems, stochastic descriptions are commonly employed when modeling open systems. This reflects the practical necessity of coarse-graining over environmental degrees of freedom, whose unresolved influence is represented through noise and dissipation. As a result, stochastic formalisms provide an effective description of open systems interacting with environments possessing many unresolved degrees of freedom.

From this perspective, the stochastic formulation of DRT does not introduce new assumptions. Instead, the core principles of the theory remain unchanged, and the stochastic regime is a direct extension of the deterministic case. In practice, since stochasticity depends on the choice of system boundaries it is contingent upon coarse graining over unresolved degrees of freedom.

When additional degrees of freedom are explicitly incorporated, the effective dynamical description shifts accordingly.

#### 3.6.1 Stochastic DRT and Fokker–Planck Dynamics

In the stochastic regime, a dynamic resource system is characterized by a probability density *p*(*X, t*) defined over phase space *X*. Rather than evolving along a single deterministic trajectory, the system is characterized by a distribution of possible configurations whose time evolution reflects both directed energetic structure and unresolved environmental interactions.

Conservation of probability requires that the evolution of *p*(*X, t*) obey a continuity equation in phase space,

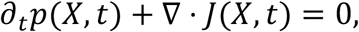

where *J*(*X, t*) denotes the probability current. This expression enforces that probability is conserved through phase space as the system evolves.

For stochastic systems, the probability current admits a standard drift–diffusion equation,

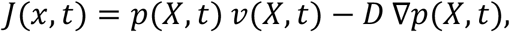

where *v*(*X, t*) is a drift velocity field and *D* is a diffusion coefficient or tensor. The drift term represents systematic transport of probability in phase space, while the diffusion term captures unbiased spreading arising from unresolved degrees of freedom. Such diffusion reflects coarse graining over environmental interactions and does not encode structured energetic organization.

Within Dynamic Resource Theory, directed stochastic transport is governed by an effective free-energy structure arising from resource allocation and resource reallocation.

The internal energy of the system is given by

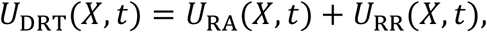

reflecting the additive contributions of allocated and reallocated energetic resources under energy conservation.

The system entropy is defined as,

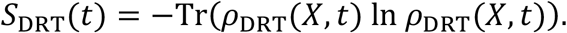

The corresponding free energy is therefore given by,

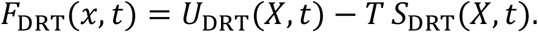

In the stochastic description, this free-energy structure induces an effective free-energy landscape *F*_DRT_(*X, t*) over phase space, whose gradients bias probability transport. Accordingly, the drift velocity is specified by a mobility-weighted gradient flow,

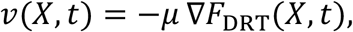

where *μ* is a mobility coefficient that converts free energy gradients into phase-space drift velocity. Substituting this form of the probability current into the continuity equation yields the Fokker–Planck equation [107] governing the time evolution of the probability density under DRT,

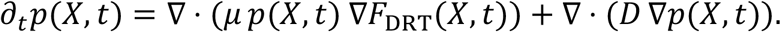

The first term (drift), *∇ ·* (*μ p ∇F*_DRT_), biases probability flow along free-energy gradients, introducing directed organization in phase space toward energetically favorable configurations. The second term (diffusion), *∇ ·* (*D ∇p*), describes stochastic spreading due to unresolved environmental coupling; it is independent of the free-energy landscape and reflects coarse-grained fluctuations rather than structured, system-level DRT interactions.

The diffusion coefficient *D* is a system- and environment-dependent parameter that sets the strength of this stochastic spreading and therefore encodes the effective stochastic dispersion of the system. It captures the net influence of unresolved degrees of freedom and interactions on probabilistic transport. In practice, *D* may be treated phenomenologically (e.g., as a constant or tensor) and estimated from observed fluctuations.

### 3.7 Interim Summary

In this section, DRT was developed as a general physical framework for describing self-organizing systems under energetic and thermodynamic constraints. RA defines the energetic structure governing stable system configurations, while resource reallocation RR provides the adaptive mechanism through which systems dynamically respond to internal fluctuations and external demands. Together, RA and RR capture the complementary roles of stability and flexibility within a single theoretical formalism.

Therefore, the allocation–reallocation structure is not an auxiliary construction but follows directly from the foundational constraints of the framework. Given Postulates 1–3, any physically admissible redistribution must correspond to a thermodynamically constrained modification of the energetic baseline configuration defined by RA. In this sense, RR represents the minimal structure required to reconcile Hamiltonian time evolution with thermodynamic irreversibility.

The evolution of dynamic resource systems is therefore constrained by fundamental variational and thermodynamic principles, ensuring physical consistency across deterministic, variational, and stochastic regimes. By remaining agnostic to specific dynamical forms and treating the allocation–reallocation mechanism as invariant, DRT establishes the general conditions under which self-organization is physically admissible.

## 4. Neural Resource Theory: DRT in Neural Systems

### 4.1 From DRT to Neural Systems

Dynamic Resource Theory (DRT) provides a general physical framework for systems that allocate and dynamically reallocate internal resources under energetic and entropic constraints. In this section, we specialize this framework to biological neural systems, referred to as Neural Resource Theory (NRT).

The nervous system is an especially natural substrate for DRT. Neural systems are energetically costly, tightly constrained by metabolism, and often required to allocate and reallocate resources across space and time in response to internal demands and external perturbations. A neural system’s physiological resources, including metabolic energy in the form of ATP, electrochemical gradients across membranes, neuromodulatory support, and glial-mediated regulation provide the physical substrate that enables neuronal interactions to occur and be maintained over time.

When DRT is applied to neural systems, the general notions of resource allocation (RA) and resource reallocation (RR) acquire a concrete physiological interpretation as neural resource allocation (NRA) and neural resource reallocation (NRR). The corresponding Hamiltonian takes the form

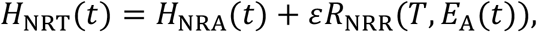

where *H*_NRA_ denotes baseline neural resource allocation and *εR*_NRR_ captures dynamic reallocation dependent on available neural resources. From the perspective of NRT, the functional consequence of these physiological resources is expressed through the interaction structure of the nervous system itself. Neural activity arises through synaptic transmission, electrical coupling via gap junctions, and coordinated population-level interactions, all of which are energetically supported and constrained by the underlying resource supply. NRT therefore treats neural resources not as isolated biochemical quantities, but in terms of the interaction capacities they collectively sustain, through which neural systems persist and remain viable.

Representative examples of neural resources and their allocation and reallocation are summarized in Table 1.

**Table 1.** Neural resources and their allocation (NRA) vs reallocation (NRR)

| Neural resource | NRA (allocation) | NRR (reallocation) |
| --- | --- | --- |
| **Blood flow / perfusion** | Sustains metabolic support to neural tissue | Task-evoked redistribution; regional perfusion shifts |
| **Oxygen & glucose** | Enable oxidative metabolism and ATP production | Increased uptake in task-relevant circuits |
| **ATP availability** | Sets energetic ceiling for neural work and interaction | Transient diversion toward coordinated or high-demand activity |
| **Ionic gradients (Na<sup>+</sup>, K<sup>+</sup>, Ca<sup>2+</sup>, Cl<sup>-</sup>)** | Maintain excitability, inhibition, and signaling readiness | Local depletion and recovery patterns during activity |
| **Excitatory synaptic transmission (glutamatergic)** | Baseline excitatory interaction strength | Rapid up/down-regulation of excitation during task engagement |
| **Inhibitory synaptic transmission (GABAergic)** | Baseline inhibitory control and stability | Dynamic reshaping of excitation–inhibition balance |
| **Gap junction coupling** | Electrical continuity between coupled neurons | Dynamic modulation of synchrony and population coupling |
| **Neuromodulatory tone (dopamine, norepinephrine, serotonin)** | Sets global gain, arousal, and operating regime | Context-dependent biasing of circuit recruitment and plasticity |
| **Structural connectivity** | Long-term constraint on possible interactions | Slowly refined via plasticity (NRA → NRA') |

Advances in neuroscience have already produced successful descriptions of neural dynamics across multiple levels of organization, including conductance-based formulations of single-neuron behavior, population-level rate models, and spatially extended field descriptions of large-scale brain activity. Neural Resource Theory unifies these descriptions within a single physical framework, while remaining fully compatible with established neuroscience. From the perspective of NRT, resource-constrained neural dynamics unfold within a landscape defined by structural, metabolic, and homeostatic constraints.

Changes in neural interaction strength and coordination are well known to occur through established neuroplastic mechanisms. These include Hebbian forms of plasticity, in which coordinated activity strengthens effective coupling between neurons, as well as synaptic pruning, whereby underutilized connections are selectively removed, releasing energetic resources that may be reallocated to support more functionally relevant networks. In this way, Neural Resource Theory provides a physically grounded framework for recasting neural organization and plasticity in terms of energy allocation and reallocation, while remaining fully aligned with established findings in neuroscience.

### 4.2 Neural Resource Theory

Neural Resource Theory provides a physical description of how energetic constraints regulate and enable interactions among neural elements. Inheriting its formal structure from DRT, NRT does not replace existing neural models or introduce new equations of motion for neural activity. Rather, it specifies how finite energetic availability shapes and modulates neural dynamics through its influence on the strength and structure of permissible interactions among neural constituents.

From the perspective of NRT, internal neural energy is described in terms of interactions between neural elements. These elements include both neurons and glial cells, which jointly regulate neurochemical gradients, ionic homeostasis, and electrical coupling through synapses and gap junctions.

For a synaptic contact or gap junction between units *i* and *j*, the local interaction energy is written as

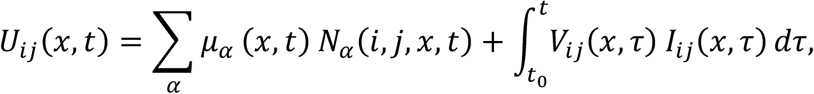

where the first term captures chemical contributions associated with neurotransmitter release, binding, uptake, and related molecular processes, and the second term captures electrical work associated with transmembrane voltage differences and currents. This expression provides a physically grounded representation of internal neural energy at the level of interacting elements and directly reflects known neurobiological mechanisms, including recent findings of a gating mechanism by which astrocytes dynamically respond to neurotransmitters and modulate neuronal circuits [108, 109].

Over mesoscopic spatial volumes and temporal windows, these microscopic interaction energies can be coarse-grained into effective resource variables and coupling parameters. In this coarse-grained description, NRT does not track individual molecular events, but instead characterizes how collections of neural interactions draw upon and redistribute energy to support circuit- and network-level activity.

System-level internal neural energy can then be calculated through spatial aggregation of these local interaction energies over the nervous system domain Ω_NS_ formed by the union between the central nervous system and the peripheral nervous system ·_CNS_ ∪ Ω_PNS_.

This systems level internal neural energy is expressed as

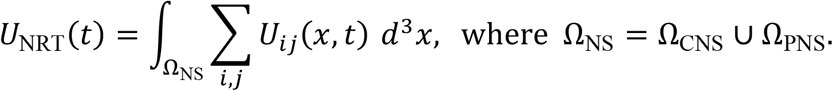

Figure 3 illustrates how system-level internal neural energy can be constructed from local interactions among neural elements across spatial scales. We begin by considering a simplified neural subnetwork composed of a small number of interacting regions, here illustrated by the thalamus (Thal), primary visual cortex (V1), and posterior parietal cortex (PPC), connected by anatomically and functionally plausible pathways (Fig. 3i).

**Figure 3.**
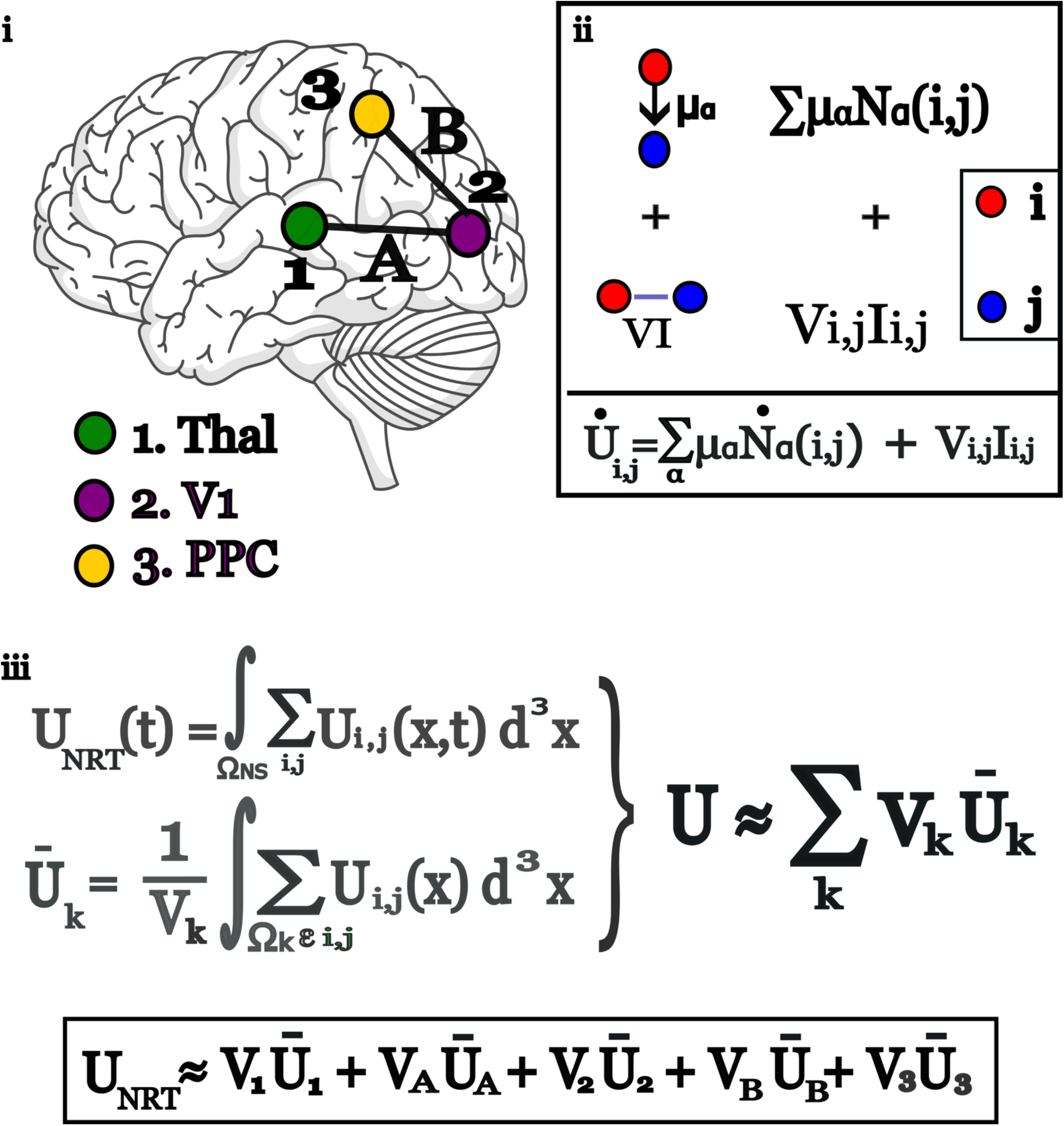
Multiscale construction of Internal Neural Energy. (i) A simplified neural subnetwork composed of interacting neural populations including the thalamus (Thal), primary visual cortex (V1), and posterior parietal cortex (PPC), connected by anatomically and functionally plausible pathways represented by A and B. (ii) Internal neural energy is defined at the level of pairwise interactions between neural elements *i*and *j*. The instantaneous interaction power is decomposed into chemical and electrical contributions. The chemical term 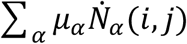 represents the rate at which neurotransmitter-mediated interactions contribute to local energetic change, where *μ*_;_ denotes the chemical potential of neurotransmitter species *α* and 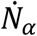 denotes the rate of engagement of those interactions. The electrical term *V*_*ij*_*I*_*ij*_ represents electrical power associated with transmembrane voltage differences *V*_*ij*_ and currents *I*_*ij*_, including synaptic and gap-junction coupling. Together these terms describe the instantaneous interaction-level power underlying neural signaling. (iii) System-level internal neural energy *U*_NRT_(*t*) is obtained by integrating local interaction energies over the nervous system domain. In practice, this may be approximated as a weighted sum of effective regional contributions, *U*_NRT_(*t*) ≈ ∑_*k*_ *V*_*k*_ *Ū*_*k*_, where *V*_*k*_ denotes the volume of region *k* and *Ū*_*k*_ its mean interaction energy density.

At the level of individual interactions, internal neural energy is defined pairwise between elements *i* and *j* through chemical and electrical interactions arising from neurotransmitter-mediated processes, ionic gradients, and transmembrane currents (Fig. 3ii). These interactions capture the physically grounded mechanisms by which neurons and glial cells support and constrain signal propagation. At larger spatial and temporal scales, these local interaction energies may be coarse-grained by aggregating across spatial volumes and interacting elements. By integrating over functionally relevant neural volumes, the full system-level internal neural energy *U*_NRT_(*t*) can be approximated as a weighted sum of effective regional contributions, providing a practical bridge between microscopic interaction energies and macroscopic neural energetics (Fig. 3iii).

Within this framework, NRA inherits the gauge theoretic structure introduced in DRT (Section 3.2) and is shaped by biological factors such as metabolic efficiency, energetic stability, and internal homeostasis. NRA describes moment-to-moment availability of energy to propagate neural signals. The system organizes its resources into a relatively stable configuration aligned with intrinsic network activity. In the human brain, this corresponds to resting-state network dynamics, where coordinated activity within systems such as the default mode network reflects the brain’s default resource distribution [110].

NRR describes the dynamic reallocation of available internal neural energy *E*_A_ to more efficiently facilitate neural interactions. In this formulation, NRA describes a relatively stable baseline configuration while NRR acts as the mechanism that perturbs and reshapes in response to demand. Over time, repeated NRR episodes progressively refine NRA, allowing the system to settle into more stable and efficient energy configurations relative to its environment.

Reallocation operates through changes in effective coupling strength, synchronization, and coordination among neurons and neural populations. In this way, NRR does not modify the form of the underlying neural equations of motion but instead modulates how strongly different elements participate in collective dynamics.

At the system level, reallocation dynamics are governed by a neural reallocation free energy defined over available resources, *E*_A_(*t*) which denotes the energy currently accessible for reallocation by NRR. It excludes energy already committed to NRA, represented by *E*_NRA_(*t*) as well as energy locked into essential maintenance functions, such as homeostatic regulation, denoted as *E*_L_ and is thus energy that is locked and is irreversibly committed to maintaining cellular integrity and structural stability and is therefore unavailable for signal propagation via inter-neuronal interactions.

The energy accessible to reallocation is thus given by

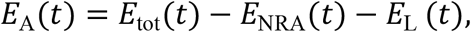

where *E*_tot_ (t) represents total neural energy at a given moment determined by the nervous system’s global metabolic budget that the body is allocating to maintain it.

The NRT system’s free energy follows directly from the DRT free energy decomposition as described in 3.4.1,

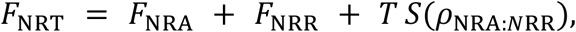

Where *F*_NRA_ is free energy of baseline neural resource allocation, *F*_NRR_ denotes the free energy associated with NRR and *TS*(*ρ*_NRA:*N*RR_), accounts for the coupling of shared energetic resources between NRA and NRR components. The reallocation function follows directly as, 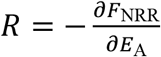.

The magnitude of the free energy change associated with neural reconfiguration is obtained by integrating the reallocation function across the range of available energy for adaptation,

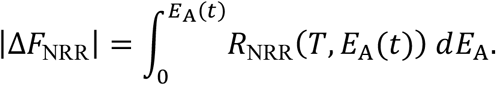

This expression quantifies the extent of energetic reallocation as the magnitude of the change in reallocation free energy.

To assess the qualitative impact of plasticity, NRT introduces the neuroplastic potential (*NP*) defined as *NP*(*t*) ≡ *F*_NRA_(*t*) − *F*_NRR_(*t*), which quantifies the net change in Helmholtz free energy resulting from reallocation. *F*_NRA_ denotes the Helmholtz free energy associated with NRA and *F*_NRR_ denotes the Helmholtz free energy associated with CRR. If *NP*(*t*) > 0, NRR results in a lower Helmholtz free energy relative to baseline NRA, indicating constructive plasticity. Conversely, if *NP*(*t*) < 0, NRR increases the Helmholtz free energy compared to NRA alone, suggesting destructive plasticity.

In neural systems, reallocation is implemented through coordination and synchronization dynamics. Changes in phase relationships among neural elements provide a mechanism by which energy is redistributed across interactions. These synchronization processes unfold on millisecond-to-second timescales [111], whereas the metabolic and vascular processes that shape allocation evolve more slowly [112].

Over longer timescales repeated patterns of activity reshape synaptic structure and circuit organization [113, 114]. From the perspective of NRT, long-term plasticity reflects gradual refinement of NRA driven by accumulated reallocation events. Established neuroplastic mechanisms, including Hebbian strengthening of coordinated interactions and synaptic pruning of underutilized connections [115] are interpreted within this framework as energy reallocation processes acting on neuronal systems.

Together, NRA and NRR provides a physically grounded implementation of Dynamic Resource Theory in nervous systems while remaining fully compatible with established models of neural activity. In the following section, we describe how this framework is realized across multiple spatial and temporal scales.

### 4.3 Multiscale Implementation of Neural Resource Dynamics

Neural Resource Theory applies across spatial and temporal scales wherever neural resources are allocated and dynamically reallocated. NRT respects established and empirically verified equations that reliably model neural activity at a given scale within their limits of uncertainty. NRT also specifies how energetic constraints shape the boundary conditions within which the dynamics being described naturally unfold. In this section, we describe NRT-informed neurophysics from microscopic neural elements through large-scale cortical organization.

#### 4.3.1 Microscale - Neurons and Local Coupling

At the microscopic scale, neural dynamics are well described by conductance-based models such as Hodgkin–Huxley formulations, which conserve charge and energy at the level of ion channel kinetics. Within NRT, these models define the local energetic constraints under which neural signaling is possible.

Ion channel conductance, membrane capacitance, firing thresholds, and refractory periods all impose energetic costs that determine when and how action potentials may be generated and propagated. These costs constitute local NRA, establishing the energetic conditions that sustain neuronal interactions. NRR at this scale is limited, transient, and highly localized, occurring through brief fluctuations in channel availability or synaptic efficacy without altering the underlying equations of motion. Microscopic neural models specify how energy is consumed during active potential generation, propagation, and synaptic transmission while NRT modulates the local conditions under which those signals occur.

#### 4.3.2 Mesoscale - Neuronal Circuits and Plasticity

At mesoscopic scales, neural interactions organize into circuits and local networks. This scale hosts many well-studied examples of neuroplastic phenomena. Hebbian strengthening reflects reallocations that concentrate energetic support toward coordinated interactions, while synaptic pruning reflects reallocations that withdraw energetic investment from underutilized connections, freeing resources for redistribution elsewhere. Neuroplastic potential, introduced in Section 4.2, provides an energetic criterion for distinguishing constructive from destructive reorganization at this level.

NRR is most naturally expressed at these scales through changes in coordination and phase synchrony. Synchronization effectively lowers the energetic cost of maintaining collective interaction patterns in a neuronal circuit by reducing redundant degrees of freedom, thereby stabilizing dynamics.

A canonical description of such coordination is provided by the Kuramoto model[29-31, 65], whose Hamiltonian can be written as

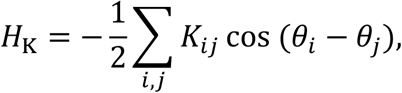

where *θ*_*i*_ denote oscillator phases and *K*_*ij*_ represent coupling strengths. Within NRT, the coupling parameters *K*_*ij*_ are interpreted as energetic investments in coordination. Increasing coupling reallocates available neural energy toward synchronized interaction, lowering the energetic cost of coherent signal propagation across neural circuits.

Here, energetic redistribution reshapes effective coupling strengths across synapses and pathways, enabling interaction patterns to evolve over longer timescales. It is common and often necessary to account for effective sources of variability arising from coarse-graining, finite population size, heterogeneous coupling, and transmission delays. Such effects are routinely represented through stochastic terms, phase noise, or distributed delays in coordination models. Since NRT inherits its formal structure from DRT, it can be implemented within stochastic frameworks, including Fokker–Planck descriptions of time evolution.

In embodied systems, coordination must support task-dependent stability and controlled transitions between interaction modes, including coordination across neural and muscular interfaces. The Haken–Kelso–Bunz (HKB) model, developed in movement science to describe rhythmic motor coordination, provides an extension of phase-based coordination dynamics into this regime[33, 67]. HKB dynamics are commonly represented through an interaction potential defining an effective stability landscape for relative phase coordination described as,

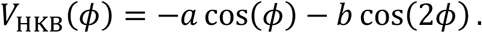

From this perspective, HKB may inform how resource reallocation can be extended beyond neuronal synchrony to incorporate neuro-muscular coordination, without introducing new principles beyond those already captured by NRT.

#### 4.3.4 Macroscale - Neuronal Populations and Field Scales

At macroscopic scales, neural activity is commonly described using neural mass models and Neural field theory (NFT). Neural mass models, such as the Wilson–Cowan framework, describe the average activity of interacting neuronal populations by capturing excitation–inhibition balance, synaptic kinetics, and population gain. NFT extends this approach by treating cortex as a continuous medium with connectivity kernels, propagation delays, and wave-like dynamics, and can be understood as a mean-field description of large-scale neural activity. These frameworks have successfully accounted for oscillatory rhythms, traveling waves, metastability, and large-scale functional organization observed in cortex.

Within NRT, neural mass and neural field models define the state dynamics of population-level neural systems. Resource allocation shapes baseline excitability, gain, and effective coupling across populations, while resource reallocation modulates these quantities dynamically by redistributing energetic support across regions and pathways. In this way, energetic constraints shape the resource landscape in which population and field dynamics unfold, without altering the mathematical form of the governing equations.

At the field level, changes in available energy influence the strength and spatial reach of coupling kernels, the stability of wave propagation, and the accessibility of distinct dynamical regimes. Transitions between functional states such as shifts between localized activity, large-scale synchronization, or transient exploratory dynamics can be interpreted as energetically mediated reallocation of population-level coordination.

This interpretation aligns with empirical evidence linking metabolism, neuromodulation, and vascular dynamics to large-scale neural activity. Rather than treating these influences as external drivers, NRT incorporates them as intrinsic components of the energetic constraints governing population-level behavior.

In this way, NRT complements neural mass and neural field models by providing a physically grounded account of how energetic availability shapes large-scale neural dynamics, while preserving their established mathematical structure.

### 4.4 Physiological Grounding and Observable Measures in Neural Resource Theory

#### 4.4.1 Experimental Modalities

Within NRT, resource allocation and relocation correspond to observable features of neural activity that are already routinely measured in laboratory and clinical settings across spatial and temporal scales. Mapping NRT to observables across experimental modalities emphasizes a broad approach that utilizes known tools and techniques native to each modality to best leverage their inherent spatial and temporal resolution. In Table 2, we outline how common neurophysiological and neuroimaging measures can be mapped to NRT-specific quantities that remain grounded in established neuroscience.

**Table 2.** Experimental Modalities and NRT-Associated Physiological Observables.

| **Modality** | **Physiological signal** | **NRA (allocation) observables** | **NRR (reallocation) observables** |
| --- | --- | --- | --- |
| **EEG / MEG** | Postsynaptic currents; population synchrony | Resting-state spectral structure; baseline oscillatory power; stable coherence patterns; excitation–inhibition balance | Task-evoked desynchronization/resynchronization; transient phase synchrony; changes in coherence; phase–amplitude coupling; directed influence shifts |
| **iEEG (ECoG / sEEG / DBS-LFP)** | Local field potentials; synaptic and transmembrane currents | Stable local rhythms; baseline coupling strength; persistent PAC structure | Rapid, spatially precise reallocation dynamics; burst synchronization; transient directional coupling; causal perturbation responses |
| **fMRI (BOLD)** | Hemodynamics linked to metabolic demand; relative deoxygenated hemoglobin concentration. | Resting-state networks; baseline functional organization; default resource distribution | Task-evoked activation; dynamic functional connectivity; state transitions; network reconfiguration |
| **Perfusion MRI (ASL)** | Cerebral blood flow | Regional energetic supply set-point | Task-dependent redistribution of perfusion; sustained reallocation costs |
| **sMRI / dMRI** | Structural anatomy and white-matter organization | Long-term constraints on allocation (connectome, cortical thickness) | Slow refinement of NRA through plasticity ($RA \rightarrow RA'$) |
| **MRS** | Neurochemical concentrations (e.g., glutamate, GABA, lactate, ATP proxies) | Baseline energetic and excitatory/inhibitory constraints | Shifts in metabolite ratios reflecting sustained reallocation |
| **fNIRS** | Cortical hemodynamics | Baseline oxygenation patterns | Task-related redistribution during naturalistic behavior |
| **FDG-PET** | Glucose uptake | Global and regional metabolic baseline; long-timescale NRA | Integrated energetic demand associated with prolonged task engagement |

For example, in fMRI, NRA in the human brain would naturally be mapped onto resting-state network dynamics measured through BOLD signal fluctuations [116]. Coordinated activity within large-scale systems such as the default mode and dorsal attention networks reflects the brain’s baseline resource distribution in the absence of task-induced strain [110]. Evidence of reallocation of neuronal resources would be reflected in changes in synchrony measures such as functional connectivity [117] and functional network connectivity [118], as well as changes in directed connectivity measures such as dynamic causal modeling, structural equation modeling, Granger causality, and transfer entropy [119, 120]. Imposing constraints imposed by structural data further improve interpretability enabling estimations in changes in effective connectivity [121, 122].

#### 4.4.2 Predictive Validity of NRT: Sleep–Wake Energetics and Neuroplasticity

To provide empirical grounding for NRT, we examine the canonical neural contrast between sleep and wakefulness.

The formal structure of NRT is captured by

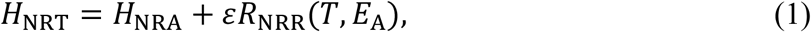

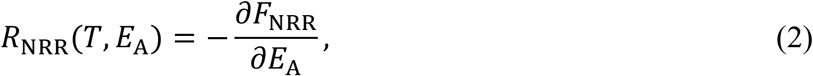

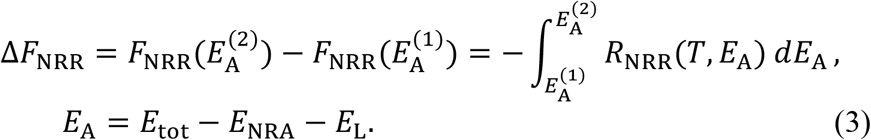

Equation (1) defines neural system energy expenditure as the sum of baseline allocation dynamics *H*_NRA_ and reallocation contributions governed by *R*_NRR_. Equation (2) links the reallocation rate to the gradient of a free energy function *F*_NRR_, introducing a thermodynamic landscape that governs reallocation efficiency. The integral form above follows directly from Eq. (2) and expresses the change in free energy across a finite interval of available energy. Equation (3) defines the energy pool available for NRR after accounting for baseline neural allocation and biologically locked energy such as homeostatic maintenance and vital functions.

From Eq. (3), differences in baseline allocation between behavioral states imply differences in available energy for reallocation. If 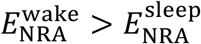, then for fixed *E*_tot_ and *E*_L_, 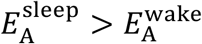.

Since NRR evolves according to gradient descent of *F*_NRR_ along *E*_A_, an increase in available energy expands the states energetically accessible to NRR. Under this condition, the magnitude of the free energy change associated with NRR transitions is greater during sleep than during wakefulness,

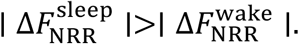

This result predicts that reduced baseline energy expenditure increases the thermodynamic capacity for dynamic reallocation, thereby enabling greater plastic reconfiguration during sleep. Empirical findings report that brain energy expenditure during non-rapid eye movement sleep decreases to approximately 85 percent of its waking value [123, 124]. This aligns with

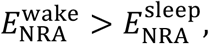

since *E*_NRA_ is the baseline internal allocation energy. A reduction in baseline expenditure therefore increases the energy available for reallocation.

Consistent with this prediction, NREM sleep has been repeatedly associated with enhanced neural plasticity relative to wakefulness [125, 126],

Within NRT, this corresponds to a larger free energy excursion along the NRR landscape during sleep, facilitating neuroplastic adaptations that improve overall system configuration in response to energetic strain or prior perturbation.

This canonical sleep–wake contrast provides a concrete neuronal instantiation of NRT by mapping measurable metabolic and electrophysiological phenomena onto the allocation– reallocation formalism of DRT. In Figure 4, we illustrate how perturbation-driven reallocation during wakefulness, relaxation during NREM, and allocation updating during REM, where metabolic expenditure is comparable to wakefulness [123, 127], collectively refine baseline neural allocation across sleep cycles. This refinement unfolds within the canonical NREM–REM sleep-stage cycle [128], situating our prediction within the broader RA–RR mechanism introduced in Section 3.3.

**Figure 4.**
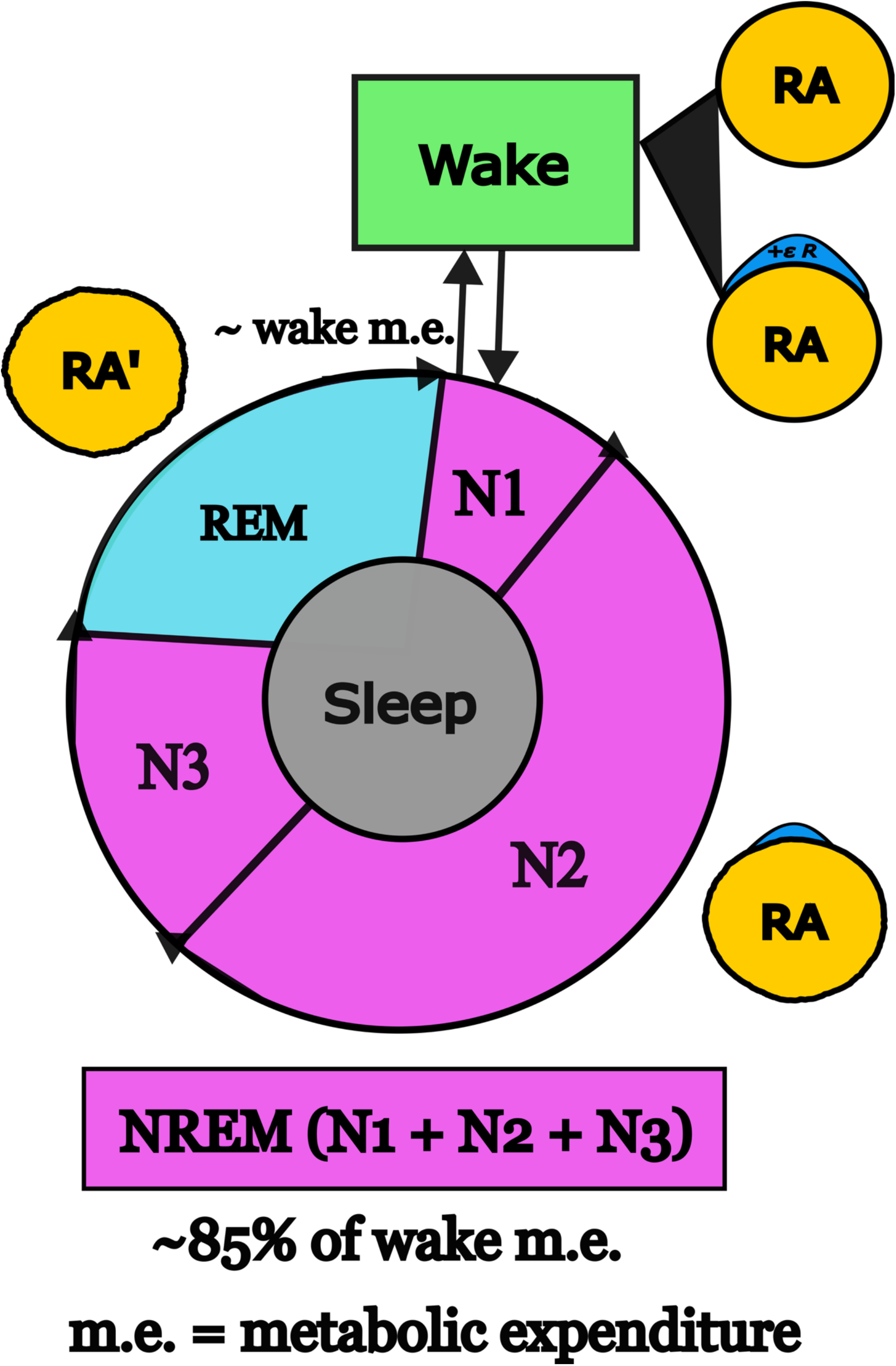
Sleep–Wake Cycle and Neural Resource Theory. Canonical sleep–wake cycle interpreted through the allocation–reallocation (RA-RR) framework of NRT. During wakefulness, neural resource allocation (NRA) sustains ongoing interaction demands while environmental perturbations transiently engage neural resource reallocation (NRR) reallocation dynamics, corresponding to the +*εR* contribution that modulates NRA. Across the day, repeated perturbation–NRR events strain NRA. Entering non-rapid eye movement (NREM) sleep is associated with decreased energetic expenditure [123], increasing the change in free energy available for reallocation 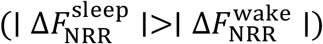. Under diminished external drive, accumulated perturbations can relax, permitting favorable conditions for the refinement of NRA. Rapid eye movement (REM) sleep intermittently elevates metabolic activity to levels comparable to wakefulness and within NRT, is interpreted as facilitating allocation updating (RA → RA’).

### 4.5 Interim Summary

This section introduced Neural Resource Theory (NRT) as a principled specialization of Dynamic Resource Theory (DRT) applied to biological nervous systems. NRT provides a physically grounded framework for understanding how energetic and entropic constraints shape neural self-organization and system adaptability through the allocation and dynamic reallocation of internal resources, while remaining fully compatible with established models of neural activity.

Neural resources were formalized in terms of interaction energies supported by biochemical, electrical, and structural substrates, and these microscopic contributions were shown to admit systematic coarse-graining into circuit-, population-, and field-level descriptions. Within this framework, NRA defines the system’s baseline energetic configuration, while NRR captures the dynamic redistribution of available energy that supports coordination, synchronization, and adaptive reorganization in neuronal systems.

NRT embeds itself across multiple spatial and temporal scales. At the microscale, energetic constraints govern local excitability and signal transmission. At the mesoscale, reallocation manifests through coordination and synchronization dynamics that underlie plasticity and circuit reorganization. At the macroscale, NRT complements neural mass and neural field models by interpreting large-scale functional dynamics as unfolding within an energetically constrained resource landscape shaped by physically grounded energetic and entropic limits.

The core quantities of NRT map directly onto observable neural signals across widely used experimental modalities, including electrophysiology, neuroimaging, and metabolic measurements. These mappings establish that NRT is empirically accessible and can be operationalized using existing data and analysis techniques.

Predictive validity was illustrated through the sleep–wake transition. NRT predicts an increased capacity for neural reconfiguration during sleep due to reduced baseline energy expenditure, consistent with well-established empirical findings on sleep-dependent plasticity.

Together, these results establish NRT as a unifying, physically grounded framework for interpreting neural dynamics, plasticity, and large-scale organization in terms of energy allocation and reallocation. In the following section, this framework is extended to Cognitive Resource Theory (CRT), where cognition is modeled as a physical, partitioned subset of NRT, applied to task engagement, cognitive effort, and behavioral performance.

## 5. Cognitive Resource Theory

### 5.1 Cognitive Resource Theory as a subset of NRT

Cognition is conventionally defined as the set of mental processes involved in knowing and awareness, including perception, memory, reasoning, judgment, imagination, and problem solving. Resource allocation [129-132] and adaptive reallocation [133-135] have precedent in cognitive neuroscience literature. The conceptualization of resource reallocation as a potential unifying principle for self-organization in complex systems first emerged from empirical studies of action video game play and visuomotor decision-making.

Cognitive Resource Reallocation (CRR) was first formally described as a hypothesis that offered a parsimonious explanation for significant shifts in brain connectivity data that correlated with the behavioral advantage observed in action video game players, who exhibited significantly faster visuomotor response times compared to age-matched healthy adults, without loss of accuracy [136, 137]. Because cognition is abstracted from the matter and energy required to construct and sustain its neural processes, it follows that these observations lent themselves to interpretation in terms of resource reallocation independent of specific sensory or motor instantiations.

Cognitive Resource Theory (CRT) provides a physical scaffolding for cognition. Rather than redefining cognitive processes themselves, CRT characterizes how neural resources and energy are dedicated to supporting the set of functions collectively referred to as cognition. CRT internal energy is treated as a subset of internal neural energy, as described in Section 4.3. Any vector field governing neural resource flow would, by necessity, project a real and meaningful component onto the cognitive subspace defined by the set of neural interactions supporting task-relevant cognitive processes.

CRT is formulated as a specialization of Neural Resource Theory (NRT), restricted to the subset of neural resources available for cognitive function. In this view, cognition does not introduce a new class of physical resources. Instead, it draws from a functionally defined subspace of the broader neural resource landscape, comprising neural energy that can be dedicated to supporting cognitive processes under varying task demands. From the perspective of CRT, cognition is thus defined as the set of neural interactions that collectively support processes involved in knowing and awareness. By grounding cognition in the dedication and redistribution of neural resources, CRT provides a principled physical framework for understanding the constraints imposed on the mechanisms that support and maintain cognitive faculties.

### 5.2 Cognitive Resource Allocation and Reallocation

Cognitive resources as a subset of neural resources inherit the energetic structure defined by NRT, in which internal neural energy is expressed in terms of local interactions between neural units (Section 4.2). Within this framework, CRT formalizes cognitive resource allocation (CRA) and reallocation (CRR) as dynamic energetic processes embedded in physically plausible neural dynamics, described by the Hamiltonian

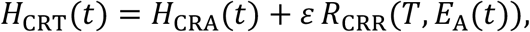

where ⟨*H*_CRA_(*t*)⟩ = *U*_CRA_(*t*) defines the internal energy associated with CRA. The quantity

*E*_A_(*t*) denotes the energy available for reallocation by CRR, and *T* is the local cognitive temperature of the CRT system. The reallocation term is defined thermodynamically as

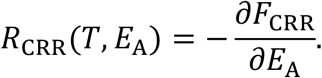

Unlike CRA, which establishes a relatively stable energetic configuration, CRR governs adaptive responses to cognitive demands. Because reallocation represents a departure from the baseline defined by CRA, its thermodynamic contribution cannot be subsumed under the free energy associated with allocation. Instead, CRR is characterized by its own free energy profile, capturing the trade-off between investing internally available energy and overcoming configurational entropy during task engagement.

The energy available for reallocation, *E*_A_(*t*), represents the portion of the cognitive system’s total energetic budget *E*_tot_(*t*) not already committed to baseline cognitive operations *E*_CRA_(*t*) or constrained by indispensable physiological requirements *E*_L_(*t*). Accordingly, *E*_A_(*t*) is given by

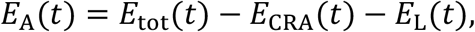

Together, CRA and CRR define the dynamic energetic landscape over which cognitive processes may occur. CRA establishes the energetic context for cognition, while CRR enables adaptive reconfiguration within that context. CRT adopts from NRT both the expression for the extent of energetic reallocation and the associated neuroplastic potential to assess its qualitative impact (Section 4.2). The magnitude of the change in free energy reallocated by CRR is given by,

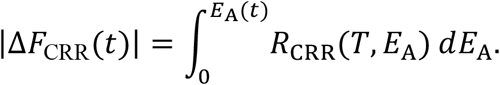

Correspondingly, neuroplastic potential is defined as, *NP*(*t*) ≡ *F*_CRA_(*t*) − *F*_CRR_(*t*), where *F*_CRA_ denotes the Helmholtz free energy associated with CRA and *F*_CRR_ denotes the Helmholtz free energy associated with CRR. If NP(*t*) > 0, CRR results in a lower Helmholtz free energy relative to baseline CRA, indicating constructive plasticity. Conversely, if NP(*t*) < 0, CRR increases the Helmholtz free energy compared to CRA alone, suggesting destructive plasticity.

### 5.3 Task-Related Energetic Expenditure in Cognitive Resource Theory

CRT is a constrained specialization of NRT, which itself derives from the DRT formalism. As established in Section 3.4, DRT admits a thermodynamic description in which system behavior is characterized by a partition function and an associated density matrix. Section 3.5 reformulates the same dynamics variationally, using action principles and path-integral weighting.

Throughout, natural units are adopted for action and inertial scale by setting the reduced Planck constant and effective mass to unity, while Boltzmann’s constant is retained to preserve the physical interpretation of temperature and energetic cost.

In CRT, cognitive load associated with a cognitive task is defined as the energetic cost associated with sustaining a task relative to CRA. Under this framework, cognitive effort may be interpreted as the energetic load required to sustain task performance during the period of its engagement.

From this perspective, effort is not identified with instantaneous energetic expenditure, but with the excess energetic cost accumulated as the system maintains a constrained cognitive configuration required for task performance.

Cognitive processes support a task-defined cognitive subspace *V*, representing the anatomical substrate actively supporting a given cognitive process at a given moment in time.

This subspace is defined as the combined volume of the engaged cortical and subcortical regions together with the white matter pathways linking them. As a concrete example, consider a visuomotor decision task requiring the transformation of visual motion information into action-relevant spatial representations. Such processes are canonically associated with dorsal stream circuitry, including occipital and parietal regions involved in motion processing and sensorimotor integration [138-140]. Regions such as the superior occipital gyrus and superior parietal lobule are regarded as task-relevant substrates supporting visuomotor transformations, consistent with their established functional roles within the dorsal visual stream [138]. In this context, *V* can be approximated as the combined volumes of these regions together with the white matter pathways connecting them. Given the physical separation between regions, long-range white matter tracts are expected to dominate interactions, with direct gray matter continuity or gap-junction coupling playing a minimal role.

The number of active cognitive units is denoted by *N*, representing the population of participating elements such as neurons, synapses, and glial components. In practice, *N* can be approximated using structural neuroimaging data by jointly considering regional tissue volumes and connectivity structure and is fixed on the time scales relevant for a given task. Gray matter volumes or voxel counts provide a coarse-grained proxy for local tissue capacity, while tractography-derived measures such as streamline density, tract volume, or connection strength offer an estimate of the connective tissue supporting task execution. This allows a principled and empirically grounded approximation of the effective system size engaged during a cognitive process. For a given momentary configuration while engaging some cognitive process, the CRT system may be treated as a dynamic equilibrium, permitting well-defined thermodynamic derivatives.

Cognitive temperature is therefore defined as a local, effective thermodynamic parameter characterizing the task-engaged subspace,

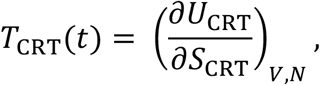

where *U* denotes internal energy and *S* the entropy of the full CRT system restricted to the task-relevant subspace.

Introducing the inverse cognitive temperature, 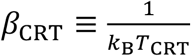, the CRA partition function is denoted *Z*_CRA_(*β*_CRT_) ≡ *Z*_0_(*β*_CRT_). This partition function characterizes the task-relevant subspace under baseline allocation, defined over the same subspace (*V, N*) and at the same cognitive temperature *T*_CRT_.

In the variational formulation, the cognitive cost functional, 𝒦_CRT_[*q*(*t*)], decomposes into baseline allocation and reallocation contributions,

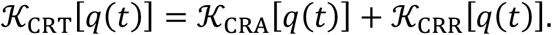

The baseline partition function is given by

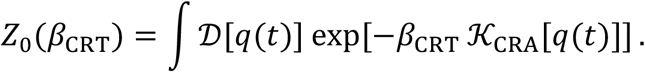

The expected baseline allocation cost is therefore

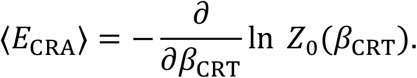

In empirical applications, ⟨*E*_CRA_⟩ may be reasonably approximated from resting-state or low-demand baseline data defined over the same task-relevant subspace. Such measurements provide an empirical proxy for CRA. Within this framework, system trajectories *q*(*t*) may be estimated using directed connectivity measures such as dynamic causal modeling or Granger causality, constrained by known anatomical pathways to define the space of feasible trajectories. Within DRT, energetic quantities associated with task performance are treated as observable functionals defined on system trajectories.

The energetic cost of sustaining a task is therefore defined as the task energy functional

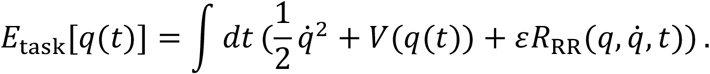

The expected energetic cost associated with task performance is computed directly from DRT as

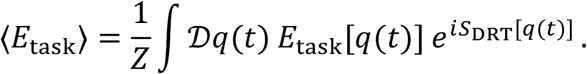

Empirically, ⟨*E*_task_⟩ corresponds to the energetic expenditure of the CRT system during task performance and may be estimated from task-evoked signal power or amplitude, such as BOLD responses in fMRI or event-related potentials and spectral power in EEG, evaluated over the task-relevant subspace.

#### 5.3.1 Cognitive Load and Efficiency

CRT frames cognitive load in terms of a thermodynamic budget. The baseline allocation ⟨*E*_CRA_⟩ parameterizes the system’s capacity to produce cognitive work ⟨*W*_out_⟩, while the task ⟨*E*_task_⟩ imposes an energetic demand ⟨*E*_demand_⟩. This provides a principled lens for interpreting resource mismatch and reallocation dynamics. The formulation parallels broader theories of dynamical optimization, including control theory, attractor dynamics, and constrained trajectory planning, where systems reconfigure themselves in response to excess or scarcity to optimize performance under changing conditions [141-143]

An energy-normalized measure of cognitive load *η*_load_ is given by the ratio between the energetic demand ⟨*E*_demand_⟩ and the baseline capacity of the system to produce useful work ⟨*W*_out_⟩, which can be described by,

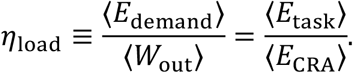

Values of *η*_load_ near unity indicate that task demands are well matched to baseline allocation capacity, whereas deviations reflect under-support or over-support of the task relative to CRA and identify regimes in which compensatory resource reallocation becomes energetically relevant to system dynamics.

When energetic load is substantially below baseline capacity (*η*_load_ < 1), CRA exceeds task demands, indicating that the task can be supported without compensatory reallocation. In this regime, CRR is not energetically required for performance, though it remains available as a mechanism for adaptation. When *η*_load_ > 1, task demands exceed baseline support, indicating energetic strain on CRA and the necessity of compensatory reallocation with an associated energetic cost to sustain performance.

A bounded measure of energetic efficiency can also be defined in accordance with standard thermodynamic and engineering principles as the ratio between the useful work ⟨*W*_out_⟩ supported by baseline resource allocation ⟨*E*_CRA_⟩ and the energetic input ⟨*E*_in_⟩ required to support the task ⟨*E*_task_⟩, yielding

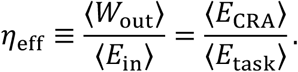

Since the task energy functional decomposes into baseline task-supporting and compensatory reallocation contributions, the total energetic input required for task execution satisfies

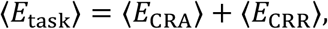

yielding the bounded thermodynamic efficiency measure defined above. Higher values of *η*_eff_ indicate that task performance is primarily supported by CRA, with minimal engagement of CRR, whereas reductions reflect increasing reliance on reallocation to compensate.

The energetic efficiency for a CRT system is therefore given by

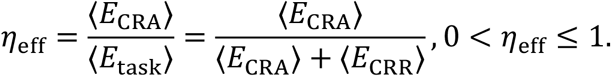

CRT does not replace existing frameworks for interpreting cognitive function, brain–behavior relationships, or task performance. Instead, it provides a physically grounded bookkeeping framework for quantifying energetic support for a cognitive process engaged during a task.

For a fixed, functionally defined cognitive process, differences in energetic cost or energetic load reflect differences in how efficiently that process is supported by the system’s baseline allocation. Enhanced performance accompanied by reduced energetic load indicates more effective baseline support, whereas increased energetic demand indicates compensatory reallocation or task strain.

To illustrate this interpretation, consider an intervention study designed to improve visuomotor decision-making performance through repeated exposure to high cognitive loads, with neural activity measured via EEG or fMRI during a left–right moving dot discrimination task[144]. This training-dependent visuomotor paradigm provides a concrete cognitive instantiation of CRT by mapping DRT’s formal structure onto task-level performance and measurable neural dynamics.

After the first phase of training, analysis reveals a behavioral difference: response times are noticeably improved without loss of accuracy. The investigators notice task relevant regions spanning thalamic input through primary motor cortex M1, which generates the behavioral output. Figure 5 depicts a simplified model characterizing the regions of interest and the anatomical white-matter connections facilitating functionally relevant rapid inter-regional signal propagation.

**Figure 5.**
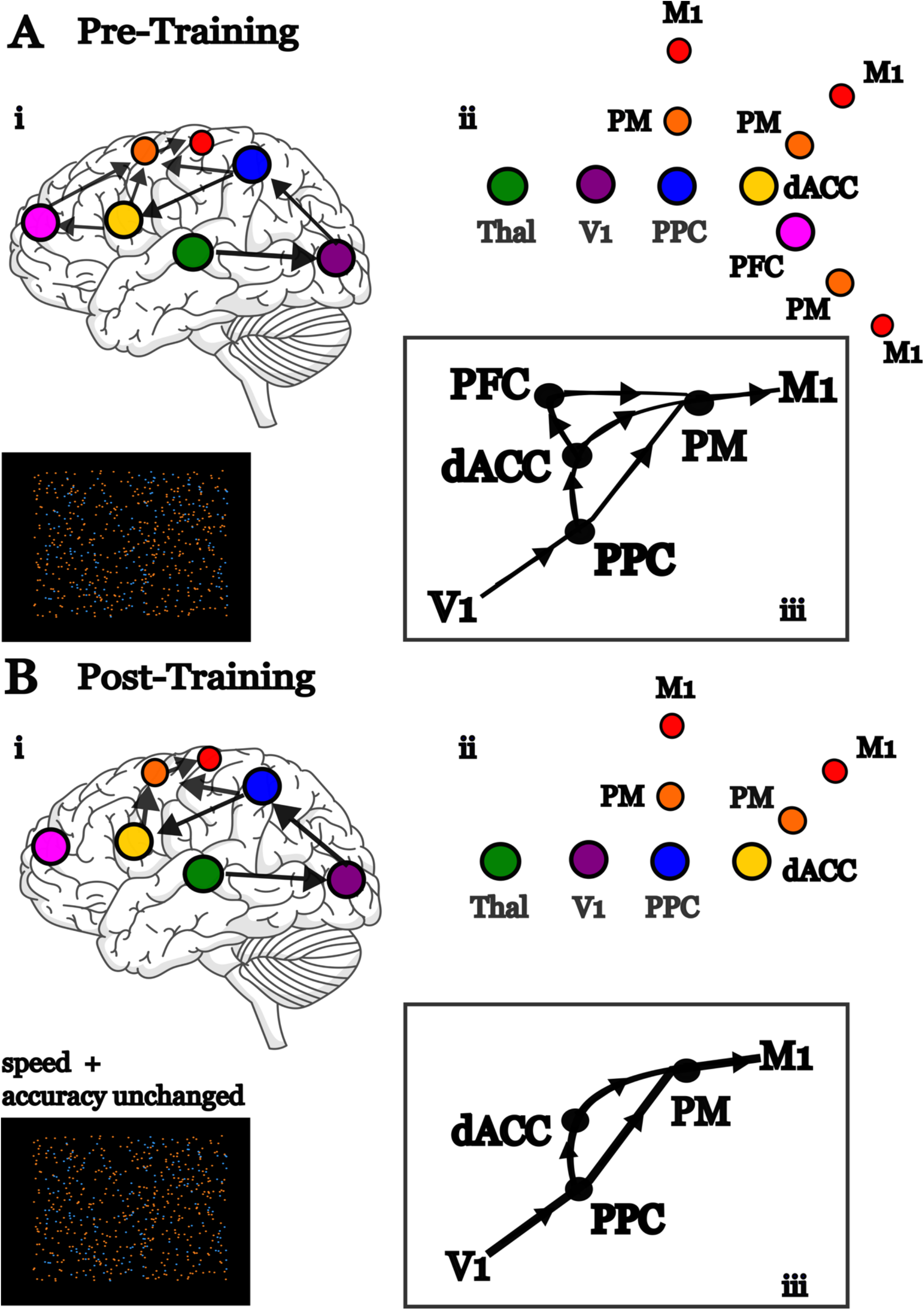
Training-dependent reconfiguration in Cognitive Resource Theory. (A) Pre-training state. (i) Visuomotor decision-making engages distributed regions including thalamus, primary visual cortex V1, posterior parietal cortex PPC, dorsal anterior cingulate dACC, pre-frontal cortex PFC, pre-motor cortex PM, and primary motor cortex M1, with prominent prefrontal modulation reflecting exertion of direct executive control. (ii) Representative routing map illustrating multiple energetically admissible interaction pathways on a 2D grid. (iii) CRT variational representation of cognitive action across these trajectories, corresponding to higher effective task energy and lower efficiency associated with pre-training. (B) Post-training state. (i) Frontal recruitment is diminished consistent with reduced demand for executive control, accompanied by faster response times with unchanged accuracy. (ii) Updated routing map showing streamlined interaction pathways following training. (iii) CRT variational representation indicating reduced but more concentrated routes, reflecting lower task energy and increased action efficiency for the same cognitive function. Within CRT, configurations of this kind admit a path-integral formulation over admissible trajectories, enabling quantitative measures of task energy, cognitive load, and efficiency.

Suppose prior to the training the group exhibits ⟨*E*_task_⟩ = 120 (arbitrary units) relative to ⟨*E*_CRA_⟩ = 100, yielding *η*_*load*_ = 1.20 and *η*_eff_ = 0.83, whereas post-training the group exhibits ⟨*E*_task_⟩ = 105 for the same baseline, yielding *η*_*load*_ = 1.05 and *η*_eff_ = 0.95. CRT interprets this reduction in energetic load and corresponding increase in efficiency as improved CRA support for the task. In both cases, the cognitive function remains unchanged (visuomotor decision making), but the energetic demands required to sustain it differ. The reallocation budget required for a given cognitive function may be estimated by measuring task-evoked energetic demand across graded difficulty levels and fitting a cost–difficulty function 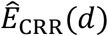. Extrapolation of this function provides an estimate of the reallocation cost required for higher-demand regimes, while task failure occurs when the required reallocation cost exceeds the state-dependent reallocation free energy *F*_CRR_, beyond which sufficient energetic support cannot be maintained.

### 5.4 Interim Summary

In this section, we established a principled framework for quantifying cognitive load, energetic cost, and efficiency within Cognitive Resource Theory (CRT). The central contribution of this formulation is not the introduction of new empirical observables, but the provision of a physically grounded interpretive layer that organizes existing measurements in terms of energetic support, reallocation demand, and system-level capacity.

Within this framework, task engagement is understood as an energetic demand imposed on the cognitive system. From the perspective of CRT, differences in task performance across individuals, groups, or can be interpreted as differences in how effectively that process is supported by baseline allocation and connectivity structure. Reduced energetic load for a fixed task reflects improved CRA support, while increased energetic demand indicates compensatory reallocation or system strain.

A key advantage of CRT is that it provides a principled quantitative separation between energetic expenditure during task engagement and the incremental cost for its execution and sustained performance. This separation allows empirical signals such as task-evoked BOLD responses or electrophysiological power to be interpreted in a manner that distinguishes baseline allocation capacity from the required energetic cost.

The cognitive load index introduced in Section 5.3 provides a compact summary of this relationship. Values near unity indicate that task demands are well matched to resource allocation capacity. Values substantially below unity indicate that resource allocation exceeds task demands, placing the system in a regime where compensatory reallocation is not energetically required for performance, but where redistribution away from the current configuration may become favorable over longer timescales. Values above unity indicate energetic strain, identifying regimes in which compensatory reallocation is necessary to sustain performance and where failure may occur if reallocation demands exceed available free energy.

By construction, these quantities are independent any one specific cognitive function. The same formalism applies to perception, decision-making, working memory, or motor control, provided the task-relevant subspace can be defined and resource allocation reasonably approximated.

Interpretation remains necessarily tied to behavioral performance and experimental context, which CRT does not seek to replace. Rather, CRT supplies a consistent energetic accounting that complements existing frameworks by making explicit the energetic conditions under which cognitive processes are supported, strained, or infeasible.

Finally, the stochastic extension of DRT described in Section 3.6 ensures that a system governed by CRT remains well defined beyond idealized deterministic trajectories. Variability, noise, and trial-to-trial fluctuations are more readily accommodated within this setting. This allows energetic cost, efficiency, and cognitive load to be defined at the level of ensembles rather than single trajectories. Cognitive state dynamics are described by stochastic differential equations whose probability densities evolve according to a Fokker–Planck equation defined over the task-relevant subspace (*V, N*). By the Feynman–Kac formula [145], solutions of the Fokker–Planck equation [107] are formally equivalent to path-integral expectation values weighted by the same reallocation cost functional, ensuring that ⟨*E*_CRA_⟩, ⟨*E*_CRR_⟩, and *η*_eff_ remain well defined under noisy dynamics. The deterministic formulation adopted here thus represents a limiting case of the more general stochastic CRT framework.

## 6. Synthesis: Neural Resource Theory and Existing Frameworks Addressing Neuronal Complexity

The frameworks outlined in Section 2 have each contributed important insights into how neural systems exhibit complex, adaptive dynamics under constraint. In this section, we synthesize NRT with gauge-theoretic approaches, statistical physics-inspired methods, dynamical population models, and the Free Energy Principle, highlighting points of alignment and key extensions enabled by the allocation-reallocation framework.

### 6.1 NRT and Gauge-Theoretic Approaches

Gauge-theoretic approaches in neuroscience have been motivated by the desire to formalize invariances, redundancies, and consistency relations in neural descriptions using a well-established mathematical language. The core motivation is that neural systems admit multiple, formally distinct descriptions that correspond to the same macroscopic function, and that symmetry principles can be used to distinguish physically meaningful invariants from representational degrees of freedom.

A few prior efforts have explored this idea in the context of brain and neuronal systems. Gauge-theoretic formalisms have been proposed in which neural dynamics are expressed on geometric or statistical manifolds, often in close connection with variational principles and approximate Bayesian inference. In these formulations, gauge structure is used to describe how internal variables or sufficient statistics may be transported consistently across different representations, thereby formalizing robustness, degeneracy, and coordination across scales. Other work has adopted more physics-inspired constructions, including lattice-like models in which synaptic or coupling variables play roles analogous to gauge fields, with the aim of studying collective behavior, phase structure, or learning under noise.

Across these diverse instantiations, a common theme is that gauge theory serves primarily as a descriptive and organizational framework. It clarifies which transformations of neural variables leave functional outcomes unchanged and which quantities remain invariant under reparameterization. In some cases, these ideas have been linked—directly or indirectly—to broader notions of self-organization or near-critical dynamics. However, such connections typically depend on additional assumptions about driving, feedback, dissipation, or plasticity. Gauge symmetry alone specifies what transformations are admissible but does not determine how neural systems move between configurations or how adaptive regimes are dynamically selected.

In Neural Resource Theory (NRT), NRA is described in terms of energetic gradients defined over an effective field. Because physically meaningful measurements must be independent of how coordinates are chosen or representation used to describe the system, a gauge-theoretic construction is required to ensure consistency. NRT, however, extends beyond a gauge-theoretic representation alone by describing how thermodynamically mediated reallocations can perturb the energetic baseline configuration defined by resource allocation, giving rise to a new baseline configuration.

### 6.2 RT and Statistical Physics-Inspired Methods

Section 2.2 described statistical physics-inspired methods as effective, descriptive tools that address the complexity that arises in neuronal systems.

Maximum entropy models have played a central role in the analysis of collective neural dynamics by providing a statistically principled way to describe population activity under empirical constraints. By adopting Gibbs distributions formally equivalent to Ising models, these approaches have been highly successful in capturing functional connectivity, network states, state transitions, and signatures of criticality in neural data.

NRT incorporates the pairwise MaxEnt model in its formal structure. In particular, the density-matrix formulation of NRT adopts an Ising-like representation of interacting neural elements, directly paralleling the Gibbs distributions used in maximum entropy models. In this sense, maximum entropy methods provide an important conceptual guide for the thermodynamic structure of NRT, informing how collective neural states can be represented probabilistically under constraint.

At the same time, NRT addresses several limitations inherent to maximum entropy approaches and other statistical physics–inspired frameworks. Whereas maximum entropy models are typically static and descriptive, NRT embeds itself within a dynamical resource framework. Resource availability and reallocation shape the evolution of the density matrix, allowing NRT to account for temporal dynamics, adaptation, and regime shifts without requiring ad hoc extensions or time-dependent constraints. Moreover, while prior methods have relied on effective, statistical notions of energy, NRT links these quantities to physically grounded neural resources, providing a pathway from statistical description to energetic mechanism.

### 6.3 NRT and Dynamical Population Models

Section 2.3 presented dynamical population models as a primary framework for describing non-equilibrium neural dynamics across mesoscopic and macroscopic scales. In this section, we situate NRT within these well-established dynamical population models.

Neural mass models (NMMs), such as the Wilson–Cowan equations, provide a foundational population-level description of neural dynamics by modeling the average activity of interacting excitatory and inhibitory populations. Rather than resolving individual spikes, these models describe the complexity of neuronal systems and their behavior through low-dimensional dynamical systems that exhibit oscillations, multistability, bifurcations, and regime transitions. As such, NMMs occupy an important middle ground between microscopic neural models and macroscopic measurements, and have been widely used to interpret EEG, MEG, and fMRI signals.

From the perspective of NRT, NMMs can be interpreted as effective descriptions of how collective neural activity evolves under implicit resource constraints. Parameters governing coupling strength, gain, and excitation–inhibition balance reflect how available neural resources are allocated across populations, while changes in these parameters correspond to reallocation processes that reshape system dynamics over time. Phase-based formulations, including Kuramoto-type models, are particularly natural candidates for describing NRR, as changes in phase coherence and synchrony directly reflect shifts in how energetic resources are coordinated across interacting neural elements.

Phase-based formulations of NMMs, including Kuramoto-type models, can be understood as reduced neural mass descriptions in which population activity is expressed in terms of collective phase dynamics. These formulations provide a natural bridge from lumped neural mass models to Neural field theory (NFT), where phase coherence and synchrony evolve continuously as population-level fields.

NFT provides a mean-field framework for modeling mesoscale neural dynamics by treating neural populations as spatially extended, continuous fields whose activity evolves over space and time. Rather than tracking individual neurons or spikes, NFT describes the average activity of large populations as a function of local recurrent interactions, spatial coupling kernels, and propagation delays. This approach captures a wide range of empirically observed phenomena, including oscillatory activity, traveling waves, multistability, and spatial pattern formation, and offers a link between microscopic neural interactions and macroscopic signals measured with EEG, MEG, and fMRI.

In this context, multistability refers to the coexistence of multiple stable population-level activity states under identical external conditions. These states arise from nonlinear recurrent interactions and correspond to distinct attractors of the field dynamics. Linear stability analysis about such states reveals characteristic eigenmodes, which determine the local stability properties of each attractor and the dominant spatial or temporal patterns governing the system’s intrinsic responses to small deviations.

NRT aligns closely with NFT at the population level, as both frameworks emphasize collective dynamics emerging from interactions among many neural units rather than from individual spikes. In NRT, local resource allocation and reallocation across interacting neural elements give rise to large-scale dynamical patterns that are consistent with those described by neural field models when microscopic detail is averaged out. From this perspective, NFT equations can be interpreted as effective, coarse-grained descriptions of underlying resource-mediated neural dynamics.

In this context, eigenmodes of the linearized field dynamics are frequently associated with Laplacian or Helmholtz operators shaped by cortical geometry. These eigenmodes define the characteristic spatial and temporal patterns through which population activity evolves, providing a basis for describing oscillatory activity, traveling waves, and stable or metastable patterns observed in neural data, and explaining how complex population dynamics can emerge from relatively simple interaction rules constrained by anatomy.

Both NFT and NRT naturally accommodate multistability, understood here as the coexistence of multiple stable or metastable dynamical regimes under identical external conditions. In neural field models, multistability arises from nonlinear interactions and recurrent coupling, allowing the system to occupy different attractor states. Within NRT, these attractor regimes correspond to distinct patterns of allocated resources across neural populations, with transitions between them driven by shifts in internal resource distribution or external perturbations. In this formulation, resource allocation dynamics are governed by an effective resource field introduced within the broader DRT framework, providing a continuous, Lorenz-gauge–constrained field-level description of how energy and resources are allocated and dynamically reallocated.

Plasticity can also be incorporated within both frameworks. NFT models typically represent plasticity through slow changes in coupling kernels, gain functions, or synaptic efficacy, capturing how population dynamics evolve over longer timescales. NRT complements this view by providing an energetic interpretation of such changes, describing plasticity as the consequence of progressive allocation and dynamically reallocation of neural resources resulting in modifications to the neural system’s energetic baseline configuration through updates to resource allocation.

In sum, NRT complements NFT by providing an energetic substrate on which field-level dynamics unfold. While neural field models characterize how population activity propagates through space and time, the energetic resources supporting that activity are typically implicit. NRT makes energy availability explicit through the allocation-reallocation framework, allowing mesoscale field dynamics to be interpreted directly in relation to underlying energetic constraints. This perspective aligns with empirical observations from EEG, MEG, and fMRI, where large-scale neural activity patterns reflect both dynamical organization and resource-dependent modulation, without requiring a reinterpretation of neural field equations themselves.

### 6.4 NRT and the Free Energy Principle

The Free Energy Principle (FEP) offers a unifying variational account of perception, action, and learning, framing neural dynamics as the minimization of variational free energy under a generative model of the environment. NRT shares with the FEP a commitment to variational principles and least-action formulations, and both frameworks interpret neural dynamics as gradient flows on an objective functional.

Under the FEP, Markov blankets define system boundaries and enable internal states to perform inference about external causes through sensory and active states. This information-theoretic structure provides a principled account of how neural systems can support inference under uncertainty. NRT is fully compatible with this perspective and can be understood as specifying the energetic substrate that underwrites such inferential processes.

At the same time, NRT directly addresses an open question in the FEP literature, namely the thermodynamic cost of belief updating. While the FEP formalizes inference in informational terms, recent work has emphasized that belief updating must incur a nontrivial physical cost. Modifying or discarding components of a generative model entails information erasure, which is subject to a fundamental lower bound on energy dissipation given by Landauer’s principle. By linking belief updating to neuronal resource consumption, dissipation, and reallocation toward more stable configurations, NRT provides a consistent thermodynamic extension of FEP

This connection becomes particularly clear when considering Cognitive Resource Theory (CRT) as a specialization of NRT. Within CRT, variational principles analogous to those employed in the FEP are applied specifically to task-engaged cognitive processes, where inference, attention, and decision-making are constrained by a well-defined subset of neural resources. From this perspective, CRT can be viewed as an instantiation of variational free energy minimization operating within the energetic constraints defined by NRT, clarifying how information-theoretic inference at the cognitive level is supported by physical resource allocation at the neural level.

In this way, FEP, NRT, and CRT operate at complementary levels of description. FEP provides a general variational account of inference under uncertainty, including within neuronal systems; NRT specifies the energetic and thermodynamic infrastructure required for its physical realization in neural systems; and CRT formalizes the energetic costs and constraints governing cognitive action under task demands. While this comparison focuses on neural and cognitive systems, it is important to note that, like FEP, DRT specifies a general variational framework that is not restricted to these domains and is intended to apply to resource-constrained self-organizing systems more broadly.

## 7. Implications and Future Directions

### 7.1 Future Developments for DRT

#### 7.1.1 Extending DRT Mechanics: Euler–Lagrange and Higher Order Perturbations

The Euler–Lagrange (EL) equations provide the standard route for deriving equations of motion from a variational principle and apply broadly to dynamical systems evolving under constraints [146]. Within DRT, the EL formalism describes how a system’s internal configuration *q*(*t*) evolves through its high-dimensional state space while balancing energetic costs, internal stability, and environmental demands. This framing treats self-organizing systems as constrained dynamical systems whose trajectories are shaped by both structural limitations imposed by RA and dynamic reconfiguration mediated by RR.

In DRT, system dynamics are governed by a constrained variational principle with both configuration-space (holonomic) constraints and velocity-dependent (nonholonomic) constraints. Holonomic constraints, which depend only on the system configuration *q* (and possibly time), encode architectural, anatomical, or capacity limits imposed by resource allocation. These constraints are handled in the standard way via Lagrange multipliers. Nonholonomic constraints arise from velocity-dependent effects associated with dissipation, reconfiguration costs, and irreversible resource flow, and are incorporated through a dissipation function.

The resulting Euler–Lagrange equations take the generalized form

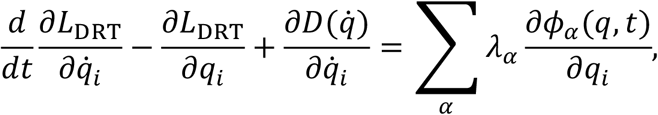

where 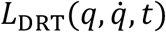 is the DRT Lagrangian, *ϕ*_*α*_(*q, t*) = 0 denote holonomic constraints, *λ*_*α*_are the associated multipliers, and 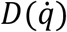 is a Rayleigh-type dissipation function capturing nonholonomic effects.

Dissipation in DRT represents the energetic cost of reallocating resources across subsystems as the system traverses its configuration space. While the lowest-order case corresponds to quadratic dissipation (Rayleigh dissipation),

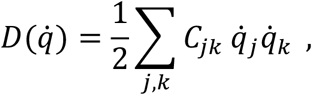

with *C*_*jk*_ encoding coupling strengths or reconfiguration costs between components. Extending dissipation beyond Rayleigh admits more general nonlinear functions of the form,

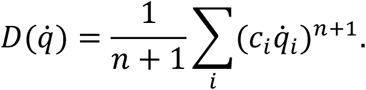

Such generalizations provide a controlled method for modeling threshold effects, nonlinear resistance, and history-dependent reallocation costs without altering the underlying variational structure. Within this framework, deviations from minimal-action trajectories arise when dissipative costs exceed the energetic capacity available under RA. In DRT terms, this imbalance signals RR engagement, which acts to redirect the system toward configurations that better align energetic expenditure with available resources. Formally, reallocation dynamics can be expressed as perturbative corrections to the RA-governed baseline, yielding a systematic expansion of the equations of motion consistent with standard perturbation theory.

Building on EL implementation, higher order RR perturbations may be required for a complete description of DRT systems. The variational structure of DRT ensures that higher-order corrections can be incorporated in a principled and dimensionally consistent manner. These effects correspond to higher-order terms in an energy-based perturbative expansion and do not require modification of the core allocation–reallocation mechanics. This may be expressed as a controlled perturbation of RA-RR dynamics. Let *L*_0_ denote the unperturbed RA Lagrangian. Reallocation enters as an energetic deformation parameterized by an energy-scale *ε*, yielding the formal expansion

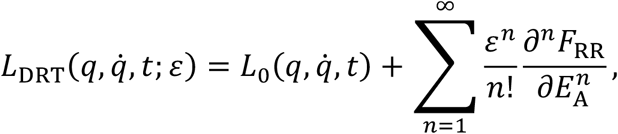

where *F*_RR_(*E*_A_) is the reallocation free energy defined with respect to the available-energy coordinate *E*_A_. Although *ε*^*n*^ carries units of energy J^*n*^, the *n*th derivative 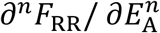 carries units of energy J^−(*n*–1)^. This structure demonstrates that DRT defines a systematic and dimensionally consistent perturbative extension of variational mechanics, and non-trivial higher-order effects may be retained as required by the system under study. As such, the Euler–Lagrange formulation provides a unified mechanical foundation for both resource allocation and adaptive reallocation dynamics.

#### 7.1.2 Extending DRT Thermodynamics

Dynamic Resource Theory is independent of the specific thermodynamic regime of the system under study. Its foundational structure depends only on its postulates which establish the existence of resource allocation and reallocation under thermodynamic constraint. As a result, DRT must be compatible with the most general thermodynamic descriptions, while also reducing consistently to stricter regimes under appropriate assumptions.

Within this general framework, DRT systems require the selection of a thermodynamic representation that specifies which state variables are treated as fixed and which are allowed to fluctuate. Thermodynamic potentials encode boundary conditions and constraint structure given by their state variables in which they are defined without altering the core structure of allocation–reallocation mechanism. This corresponds to different regime choices applied to the same physical system and are related through standard Legendre transformations.

The Helmholtz free energy provides the natural starting point for this construction and has been used throughout Sections 3-5 to construct DRT and its applications in neuronal systems and is defined as

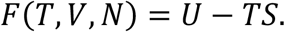

The Helmholtz free energy isolates the internal energetic degrees of freedom of a system under fixed temperature, volume, and particle number. In the context of DRT, this corresponds to the minimal thermodynamic description of internal resource allocation and reallocation, prior to introducing additional assumptions about environmental exchange, mechanical work, or compositional variability.

Other standard thermodynamic potentials arise by choosing different sets of macroscopic state variables to be held fixed or quasistatic, with the remaining variables allowed to fluctuate. Each choice corresponds to a different thermodynamic representation of the same underlying system and is characterized by a distinct set of natural variables defining the relevant potential. When pressure rather than volume is fixed, the Gibbs free energy is obtained by adding a pressure– volume contribution,

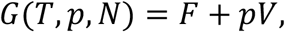

shifting the natural variables from (*T, V, N*) to (*T, p, N*). This form is particularly relevant for chemical and biochemical systems operating under fixed pressure conditions.

Similarly, the enthalpy may be expressed directly in terms of the Helmholtz free energy as

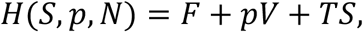

corresponding to a regime in which entropy and pressure are treated as the natural variables. Within DRT, this represents an additional constraint on to the same underlying allocation-reallocation dynamics.

When particle number is allowed to fluctuate, as in systems exchanging matter with their environment, the appropriate thermodynamic potential is the grand potential,

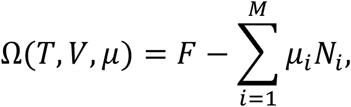

where each chemical potential *μ*_*i*_ encodes an independent particle species and imposes an additional constraint on system composition. The grand potential extends the Helmholtz description to regimes in which system composition itself becomes dynamic and provides a natural thermodynamic representation for open systems with particle exchange.

An important direction for future work is to determine which thermodynamic potential provides the most appropriate description for a given class of systems, timescales, and modes of interaction in self-organizing systems naturally suited to a DRT-based representation. Once a potential is specified, standard thermodynamic structure follows automatically, including Maxwell relations and associated stability criteria. These relations constrain response functions and admissible reallocation dynamics, providing a principled route for assessing stability, susceptibility, and robustness across thermodynamic regimes. Systematically mapping DRT behavior across these regimes represents a key step toward understanding how resource reallocation is most naturally represented in a given system of interest.

#### 7.1.3 DRT Operator Formalism

An operator formulation of Dynamic Resource Theory (DRT) provides a compact and extensible representation of resource allocation and reallocation in systems with discrete degrees of freedom, stochastic dynamics, or openness to environmental exchange. This formulation does not presume an underlying field theory or spacetime locality. Instead, reallocative dynamics are treated at the level of system states, consistent with standard operator approaches in quantum and statistical mechanics. The goal here is not to fully quantize DRT, but to demonstrate that the theory admits a clean and internally consistent operator-level description that naturally interfaces with open-system dynamics.

Under the operator formalism, the DRT Hamiltonian may be decomposed into RA and RR components as before

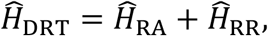

where 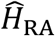 encodes baseline resource allocation and 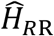 governs reallocative perturbations. Since RA structure has been specified throughout the theory, it is natural to work in the eigen-basis of 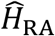, which defines the system’s preferred energetic configuration.

In this basis,

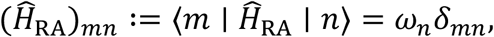

so that baseline allocation is diagonal by construction. Equivalently, the RA Hamiltonian may be written in a mode representation as

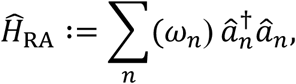

where 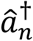 and 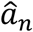 are ladder (creation and annihilation) operators that respectively raise and lower the occupation of RA-defined eigenmodes. Acting on an eigenstate of 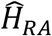, these operators increase or decrease the corresponding mode’s eigenvalue in discrete steps. The composite operator 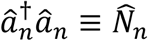 is the associated number operator, which measures the occupation of the *n*-th allocated mode.

Reallocation enters as an operator-valued perturbation of the RA-defined baseline. The reallocation Hamiltonian is written as

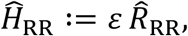

where 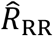 is the operator form of the reallocation function, 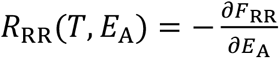, and *ε* is a quasistatic perturbation amplitude carrying units of energy and satisfying *ε*/*E*_RA_ ≪ 1. As established earlier (Section 3.3.1), *ε* scales inversely with the curvature of the reallocation free-energy landscape and controls the strength of reallocative deviations from RA-defined structure.

The operator 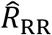 depends on the available-energy operator

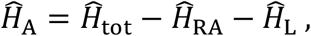

where 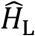 accounts for system energy that is locked and unavailable for reallocation on the timescales of interest. 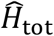 is the full Hamiltonian of the system. Although *ε* depends functionally on 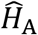, it enters only as a scalar prefactor and is therefore written without an operator hat. Since both *ε* and 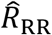 depend on 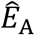, they commute.

In the RA eigenbasis, the reallocation Hamiltonian is represented by a full rank-2 operator,

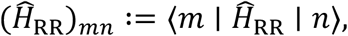

so that the DRT Hamiltonian takes the form

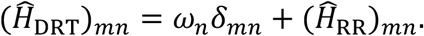

This representation makes explicit that reallocative dynamics act as controlled, energy-constrained deviations from RA-defined baseline structure. For notational convenience, the same operator may be written in ladder-operator form as

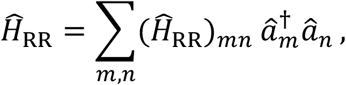

where the ladder operators provide a compact representation of the matrix elements but introduce no additional physical structure beyond that already encoded in 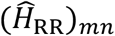.

The reallocation free energy *F*_RR_ is constructed using the von Neumann entropy,

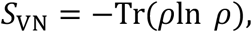

where *ρ* is the DRT density matrix. When the system explores its accessible reallocative states ergodically, *S*_VN_ reduces to the microcanonical form *k*_*B*_ln(*W*), recovering the classical definition of entropy. In classical statistical mechanics, temperature is defined through an entropy–energy relation at fixed macroscopic constraints.

In DRT, those constraints are supplied by the RA-defined configuration space, and the relevant energy variable is *E*_A_. When a scalar temperature is well defined, it is implicitly introduced through the entropy–energy dependence encoded in the reallocation function *R*_RR_. In regimes where such a temperature cannot be meaningfully defined, reallocative dynamics remain governed directly by entropy gradients without invoking thermodynamic temperature explicitly. The operator formalism accommodates both cases without modification.

When a DRT system exchanges energy, matter, or information with its environment, its state would evolve according to a Lindblad master equation,

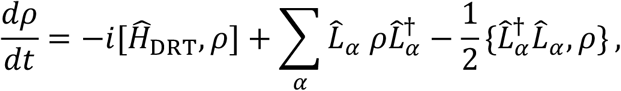

where the Lindblad operators 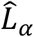 (also referred to as jump operators) describe the open, dissipative part of the dynamics, and {*a, b*} = *ab* + *ba* denotes the anticommutation relation. The structure of the Lindblad operators specifies how environmental degrees of freedom act on the system and is typically derived either from microscopic models of system–environment coupling or introduced phenomenologically. If those environmental degrees of freedom can be accounted for by expanding the system boundary to include them, the resulting combined system may be treated as closed, and the effective dissipative terms are no longer necessary.

Finally, since Lindblad evolution describes intrinsically open systems, it may be natural in some regimes to construct *F*_DRT_ using a grand canonical potential (Ω) rather than a Helmholtz free energy *F*, reflecting fluctuating system composition. In quantum or strongly correlated regimes where Boltzmann statistics are insufficient, the corresponding partition structure must be reconstructed using Bose–Einstein and/or Fermi–Dirac statistics. These changes modify the description of the system’s entropy while leaving the RA and RR Hamiltonian operator decomposition unchanged.

#### 7.1.4 Bridging Dynamic Resource Theory and the Free Energy Principle

An important future direction for Dynamic Resource Theory (DRT) lies in its relationship to the Free Energy Principle (FEP).

The FEP provides an informational variational framework describing how adaptive systems maintain their organization by minimizing variational free energy, thereby constraining perception, action, and learning through generative model optimization. DRT in its allocation-reallocation framework, complements the FEP by characterizing the mechanisms by which physical systems and their components are selectively engaged, disengaged, or reconfigured to support self-organization and functional adaptation.

From this perspective, dynamic reallocation may provide a systems-level implementation through which free energy minimization and related adaptive objectives can be realized in practice. In neural systems, this corresponds to the redistribution of energetic, computational, and network resources under changing demands. DRT places energetic efficiency at the center of adaptive dynamics by explicitly modeling the allocation and reallocation of limited energetic resources.

At the same time, DRT may capture adaptive processes involving transient network reorganization and resource constraints that are not fully specified by informational variational formulations alone.

Recalling the efficiency index constructed in Section 5.3.1, the general energetic efficiency of a DRT system is given by,

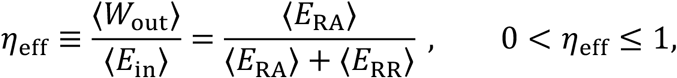

where ⟨*E*_RA_⟩ and ⟨*E*_RR_⟩, denote the expected allocation and reallocation energy contributions, respectively, and ⟨*E*_in_⟩ = ⟨*E*_RA_⟩ + ⟨*E*_RR_⟩ denotes the total energetic input supporting system dynamics. In this sense, DRT complements the FEP by providing a mechanistic account of how adaptive brain function emerges through the dynamic redistribution of limited resources. In Figure 6, we illustrate this relationship schematically.

**Figure 6.**
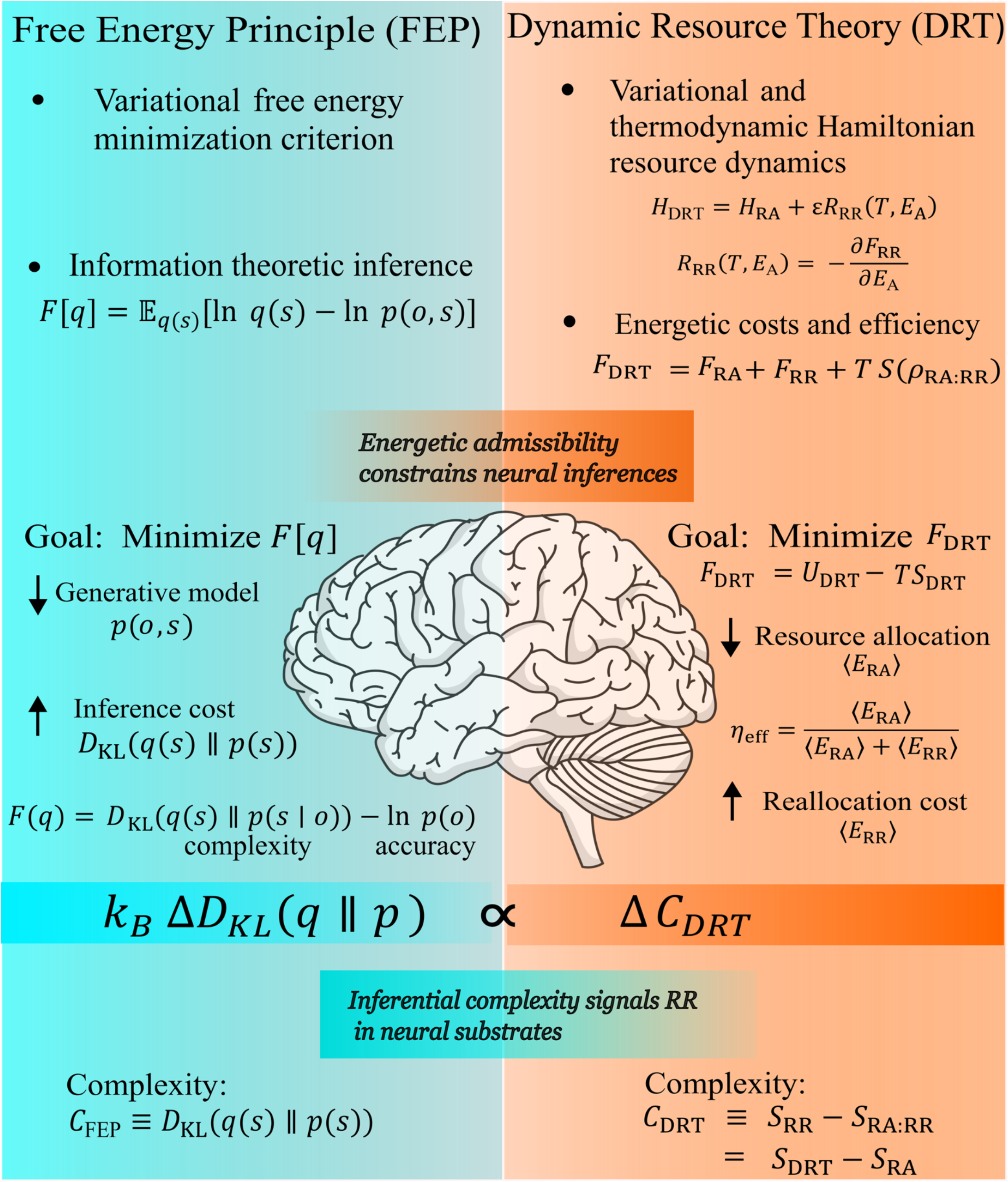
Conceptual and formal comparison between the Free Energy Principle (FEP) and Dynamic Resource Theory (DRT). The left panel summarizes FEP as a variational free energy minimization framework grounded in information-theoretic inference, where complexity is defined as the Kullback–Leibler divergence *D*_KL_(*q*(*s*) ∥ *p*(*s*)), and the objective is to minimize *F*[*q*]. In the FEP panel, top–down dynamics (↓) correspond to predictions generated by the generative model *p*(*o, s*), while bottom–up dynamics (↑) correspond to inferential complexity (inference cost) quantified by *D*_KL_.The right panel presents DRT as a thermodynamically constrained Hamiltonian framework in which baseline resource allocation *H*_RA_ is dynamically updated through resource reallocation *εR*_RR_ under energetic constraints. The corresponding system free energy is decomposed as *F*_DRT_ = *F*_RA_ + *F*_RR_ + *TS*(*ρ*_RA:RR_), with the objective to minimize *F*_DRT_. Here, *F*_RA_ denotes baseline allocation free energy, *F*_RR_ denotes reallocation free energy, and *S*(*ρ*_RA:RR_)denotes the entropy of the coupled allocation–reallocation density. In the DRT panel, allocation dynamics (↓) reflect baseline allocation energy ⟨*E*_RA_⟩, whereas reallocation dynamics (↑) reflect compensatory energetic reallocation contribution, ⟨*E*_RR_⟩ . The general energetic efficiency is 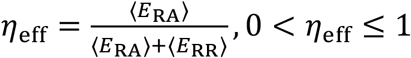, where ⟨*E*_RA_⟩ + ⟨*E*_RR_⟩ denotes the total energetic input supporting system dynamics. Complexity in DRT is defined as reallocation entropy relative to baseline allocation. The central bridge links inferential and reallocation complexity such that energetic admissibility constrains neural inference, while changes in inferential complexity signal resource reallocation dynamics.

### 7.2 Future Developments for NRT

#### 7.2.1 NRT Towards a Unified Description of Neuronal System Energy

One future direction for Neural Resource Theory is its potential to serve as a unifying description of energy in resource constrained neuronal systems across spatial, temporal, and organizational scales. As developed here, NRT provides a physically grounded framework in which baseline allocation and dynamic reallocation processes are defined independently of any specific kind of dynamical formalism, allowing it to interface with a wide range of established neural models.

From this perspective, NRT can be understood as specifying an energetic kernel that constrains and modulates neural dynamics, rather than as a replacement for existing descriptive models. Neural mass models, neural field theoretic models, and related population-level approaches describe how neural activity evolves in time and space; NRT complements these frameworks by making explicit the energetic budgets and reallocation constraints under which such dynamics occur. In this sense, NRT supplies the thermodynamic and energetic substrate upon which diverse dynamical descriptions are realized.

This embedding across scales is particularly important for linking physiological grounding to abstract dynamical models. As shown in Section 4.4, NRT-relevant quantities map onto observable measures across experimental modalities, from fast electrophysiological signals to slow metabolic and hemodynamic markers. These mappings suggest that NRT can provide a common energetic reference frame that remains consistent across EEG, MEG, fMRI, PET, and invasive recordings, without requiring modality-specific reinterpretation of core theoretical quantities.

At the mesoscale, neural mass and neural field formulations offer complementary descriptions of population dynamics, ranging from lumped excitatory–inhibitory units to spatially continuous activity fields. Within NRT, such models can be interpreted as different resolutions of the same underlying resource-mediated system, with changes in synchrony, gain, or coupling reflecting shifts in how energetic resources are allocated and dynamically redistributed. This interpretation preserves the strengths of existing models while situating them within a physically constrained energetic landscape.

More broadly, this perspective positions NRT as a scaffold for integrating neural dynamics across levels of description. Rather than privileging a single mathematical formalism, NRT emphasizes constraints that must be satisfied by any viable model of neural activity such as finite energy availability, thermodynamic dissipation, and the necessity of reallocation under strain. Future work may explore how specific dynamical frameworks such as phase-based models, field equations, or hybrid formulations can be systematically coupled to NRT-defined energetic variables, enabling principled comparisons across scales and modalities.

In this way, NRT offers a path toward unifying neuronal system energy without collapsing distinct dynamical descriptions into a single model. Instead, it provides a physically grounded reference within which diverse neural dynamics can be interpreted, compared, and constrained, setting the stage for human-specific modeling and clinical relevance addressed in the following section.

#### 7.2.2 Human-Specific NRT Models and Clinical Relevance

An important future direction for NRT lies in the development of human-specific models grounded in empirical neurophysiology. By framing neural function in terms of energy allocation and reallocation, NRT provides a physically interpretable foundation for understanding human brain dynamics across cognitive, affective, and physiological domains. This perspective has the potential to enrich existing accounts of human neurology and psychology by situating them within a unified, resource-constrained framework.

In Section 5, cognition was treated as a collection of processes that allowed Cognitive Resource Theory (CRT) to be formulated as a well-defined, task-relevant specialization of Neural Resource Theory. This framing carries a notable implication: cognitive processes, despite their complexity, can be treated as physical phenomena subject to energetic constraints, dissipation, and reallocation, like other biological systems. From this viewpoint, learning, expertise, and cognitive adaptation correspond to measurable changes in how neural resources are allocated and dynamically redistributed. Although significant work remains to translate these ideas into practice, the ability to quantify energetic costs associated with cognitive demand and adaptation on an individualized basis suggests a principled route for interpreting variability in performance, fatigue, and resilience.

Cognitive demand represents only one structured form of strain on human neural systems. At the level of NRT, quantities previously described in terms of cognitive tasks or cognitive demands are more generally treated as sustained energetic strain, stress, or load, of which task-related demand represents a special case. Other pervasive sources include intrinsic physiological stressors, affective modulation, and mixed emotional–behavioral states that are not typically described as cognitive tasks. Stress, for example, has been shown to exert heterogeneous effects depending on individual resilience and susceptibility [147], altering neural dynamics in ways that cannot be fully captured by task-based cognitive frameworks alone. NRT applies equally to these non-cognitive regimes, as they also involve shifts in baseline allocation, reallocation efficiency, and sustained cognitive load.

It is important to emphasize that within this framework, resource constraints are not inherently pathological. States commonly labeled as “stressful” or “affective” may be adaptive, context-appropriate, or even beneficial depending on environmental demands and timescales. Transient increases in energetic demand associated with fear, vigilance, or heightened arousal can support rapid action selection, learning, or survival-relevant behavior. NRT therefore does not itself assign intrinsic value judgments to neural states. Instead, strain is characterized in descriptive terms, with its functional relevance determined by context.

Accounting for such intrinsic physiological strain requires treating the nervous system as an open, non-equilibrium system. Genetic variation, pharmacological interventions, and chronic stressors would act by modifying resource allocation, altering reallocation dynamics, or imposing persistent energetic demand through ongoing exchange of energy and matter. Within NRT, these influences are interpreted as changes to system constraints rather than as distinct mechanistic categories. When open-system effects are central, the appropriate thermodynamic description shifts to a grand potential, allowing relevant chemical species and energetic resources to fluctuate without altering the core structure of the theory.

Neurodevelopment provides a particularly clear example of this open-system character. Developmental processes involve sustained energy and material exchange, changes in effective system size, and evolving constraints on connectivity and resource flow. Under these conditions, fixed-volume or closed-system descriptions are no longer appropriate. Growth, pruning, and long-timescale neuroplastic adaptation can be understood as reallocation processes unfolding under gradually changing constraints.

From a clinical perspective, this resource-theoretic viewpoint may offer a more unified account of interpreting neurological and psychiatric conditions. Although such conditions differ in symptoms and timescales, both are consistently associated with measurable alterations in neural structure, connectivity, and dynamics. Within NRT, these alterations can be interpreted as the result of constrained allocation, impaired reallocation capacity, or persistent intrinsic strain that drives neural systems toward rigid or maladaptive dynamical regimes.

Quantities related to NRA and NRR may provide interpretable biomarkers and signatures of neural system state across the lifespan. For example, from an NRT perspective it is unsurprising that recent developments have shown that the interpretability of data-driven fMRI directed-connectivity measures may be improved by incorporating structural constraints derived from diffusion MRI tractography. By restricting inferred interactions to anatomically plausible pathways, structurally constrained analyses reduce spurious directionality and link functional dynamics more directly to underlying neural architecture. In NRT terms, structural connectivity encodes baseline constraints on the flow of available resources, while directed functional interactions reflect dynamic reallocation along feasible channels.

## 8. Conclusion

The central challenge addressed here is how complexity arises in physical systems, with emphasis on resource-constrained neuronal systems. Dynamic Resource Theory (DRT) addresses the problem of complexity at the level of foundational physics. DRT identifies a minimal structural requirement: if a physical system admits an allocation–reallocation structure consistent with the postulates of the theory, then it possesses the capacity for sustained self-organization and the potential to form emergent properties. In this view, emergence does not require an additional organizing principle beyond the allocation-reallocation dynamics that define the system’s effective energetic structure. Rather, emergent behavior corresponds to self-organization at the scale where the phenomenon is naturally observed, with allocation and reallocation determining the capacity for system-level interaction.

DRT thus provides a unifying physical basis for understanding how complex behavior can be sustained and clarifies the conditions under which self-organization and emergence are physically admissible. The allocation-reallocation mechanism is treated as an invariant energetic bookkeeping that accommodates diverse dynamical descriptions by clarifying the energetic constraints and reallocation processes that makes self-organization possible. DRT concerns the minimal energetic and structural conditions under which sustained self-organization can arise in systems with finite energetic resources.

While DRT may be embedded within different physical descriptions depending on the scale of interest, it does not claim unification across fundamental forces. Instead, it specifies the conditions under which self-organization is permitted, given fundamental forces taken as ontological priors operating at their naturally occurring strengths and ranges of causal influence. In this way, DRT characterizes the constraints that govern how complex dynamics can emerge and persist across scales.

DRT establishes a foundational physical framework for resource-constrained, self-organizing systems. NRT instantiates this framework in neuronal systems by mapping its principles onto measurable observables, including electrophysiological signatures of resource redistribution, fMRI-based network energy dynamics, and metabolic energy consumption patterns. As a theory formulated at the level of general physical principles, DRT itself is not directly tested in isolation; rather, its empirical evaluation occurs through such system-specific instantiations. In this way, DRT provides the theoretical structure that enables future empirical tests and systematically derived predictions. CRT extends this structure to cognitive systems, providing a physically plausible scaffold for interpreting cognition as an energetically governed set of processes embedded within neural substrates. Thus, NRT and CRT provide domain-specific realizations and complement, rather than replace, the existing frameworks we examined.

This perspective of complexity as a physical property suggests a simple organizing constraint linking complexity to internal resource reallocation – a complexity-reallocation principle. In this view, organization arises through iterative allocation–reallocation dynamics that locally counteract configurational entropy without violating fundamental physics. The implications of this perspective extend beyond neuroscience. Since DRT is formulated independently of biological instantiation, it provides a foundation for studying emergence and self-organization across a wide range of complex systems. Future work will be required to explore additional instantiations and develop empirical methods capable of directly probing resource allocation– reallocation dynamics. Nevertheless, the framework presented here establishes a physically grounded basis for understanding how adaptive, self-organizing systems can arise and persist within an increasingly entropic universe.

## Code availability

The simulation script used to generate Figure 2 is available on the Open Science Framework (OSF) at https://osf.io/9upgw, enabling full reproducibility.

## Notes

https://osf.io/9upgw

### Competing Interest Statement

The authors have declared no competing interest.

